# Growth rate structures the dynamics of bacterial trait evolution

**DOI:** 10.64898/2026.10.22.761225

**Authors:** Chunhui Hao, Naoki Konno, Yang Zhang, Fangliang He

**Author notes:** Correspondence should be addressed to C.H. or N.K.

## Abstract

Bacterial species span orders of magnitude in growth rate, yet how its evolution is embedded within the broader evolutionary dynamics of other bacterial traits remains poorly understood. We integrated experimentally measured growth rates for 283 bacterial species with 11 genomic, metabolic, ecological and environmental traits, and used multivariate Ornstein–Uhlenbeck models to characterize how growth rate and associated traits coevolve across evolutionary time. Growth rate was best explained by models assuming distinct slow- and fast-growing evolutionary regimes, and evolved toward adaptive optima more rapidly than any other trait examined over longer evolutionary timescales, both when modelled independently and jointly with other traits. Coevolution between growth rate and associated traits was primarily structured by growth-rate regimes, suggesting growth rate serves as a pacemaker of evolution. At instantaneous timescales, six of the eight traits coevolving with growth rate showed unidirectional coupling, including all four environmental traits, with the direction varying among traits. In contrast, ribosomal RNA operon copy number and GC percentage exhibited reciprocal coupling with growth rate. These findings recast bacterial growth strategy from a correlate of trait variation to a major organizing axis shaping bacterial trait evolution across distinct evolutionary timescales.

## Introduction

Growth rate is a key bacterial life-history trait that influences how cells respond to changes in resource availability, environmental conditions and interactions with other organisms. It plays a key role in processes spanning biological scales, from pathogenesis and microbial community assembly to global biogeochemical functioning^1–11^. Over decades of study, an observation has emerged — bacterial growth rates span orders of magnitude across species^12–15^. Fast-growing bacteria such as *Escherichia coli* can double in ∼20 minutes^16^, whereas slow-growing taxa such as *Helicobacter pylori* take hours to days to complete a life cycle^17^. In extreme circumstances such as the deep subseafloor, generation times can even exceed 100 years^18,19^. This vast disparity in growth rates raises a question: how these contrasting growth strategies have evolved and been maintained across the bacterial tree of life?

Numerous hypotheses have linked bacterial growth rate evolution to traits related to genomic architecture, metabolic properties, environmental tolerance, and ecological strategy (Figure 1b; Table S1). However, empirical support for these proposed relationships has been inconsistent across studies (Table S1). For example, genome size has been hypothesised to trade off with growth rate because selection for rapid growth may favour reallocating phosphorus and nitrogen from DNA to RNA and proteins^20–22^. Yet cross-species comparative studies have reported weak, absent or context-dependent relationships between growth rate and genome size^23–25^. Similar inconsistencies arise for many other putative trait–growth relationships (Table S1).

More importantly, these hypotheses imply different coevolutionary relationships between growth rate and associated traits over macroevolutionary timescales, depending on whether evolutionary change in one trait influences, responds to, or occurs independently from change in the other (Figure 1c). However, most comparative tests, relying on traditional phylogenetic comparative approaches such as phylogenetic generalized least squares (PGLS), have focused on whether traits covary among present-day species rather than on the evolutionary processes that generated these patterns^26,27^. Similar patterns of covariance can arise when one trait influences the evolutionary response of another, when the direction of influence is reversed, when traits evolve through reciprocal coupling, or when they evolve largely independently (Figure 1c; Figure S17). Distinguishing among these alternatives therefore requires moving beyond asking which traits covary across species to understanding how their evolutionary trajectories became coupled through time, including the direction and reciprocity of these interactions (i.e., the mode of evolution) and the relative rates at which traits respond to evolutionary change (i.e., the tempo of evolution)^28^.

Multivariate Ornstein–Uhlenbeck (mvOU) model provides a way, known as the “adaptation-inertia framework”, for distinguish among alternative evolutionary hypotheses by modelling the joint evolution of continuous traits toward adaptive optima while estimating both their evolutionary coupling and the strength of selection^29–32^ (Figure 1a; Figure S9). This process-based framework allows alternative coevolutionary modes between growth rate and a focal trait (e.g., genome size) to be explicitly characterized (Figure 1c). Such OU-based phylogenetic comparative frameworks have been increasingly applied to address a wide range of macroevolutionary problems, including the coevolution of life-history traits in amniotes^33^, the joint evolution of gene expression and protein abundance^34,35^, regime-associated shifts in life-history evolution of endotherms during major habitat transitions^36^, and the inference of how diverse traits influence the macroevolution of plant growth form^37^.

Here, we investigate how the evolution of bacterial growth rate is embedded within the evolutionary dynamics of genomic, metabolic, ecological, and environmental traits. Using experimentally measured growth rates and 11 associated traits across 283 bacterial species (Figure 1b; Table S1), we applied OU-based phylogenetic comparative methods to determine whether these traits evolved independently, showed directional evolutionary dependencies, or exhibited reciprocal coupling with growth rate, while quantifying differences in their evolutionary tempos (Figure 1c). Our analyses revealed that trait coevolution is structured by contrasting growth-rate regimes. Growth rate evolves rapidly toward adaptive optima, whereas associated traits generally evolve more slowly and show distinct directional and reciprocal coupling with growth rate across evolutionary timescales. These results move beyond correlation-based approaches to provide a process-based view of how bacterial growth strategies and associated traits have evolved over macroevolutionary timescales.

**Figure 1.**
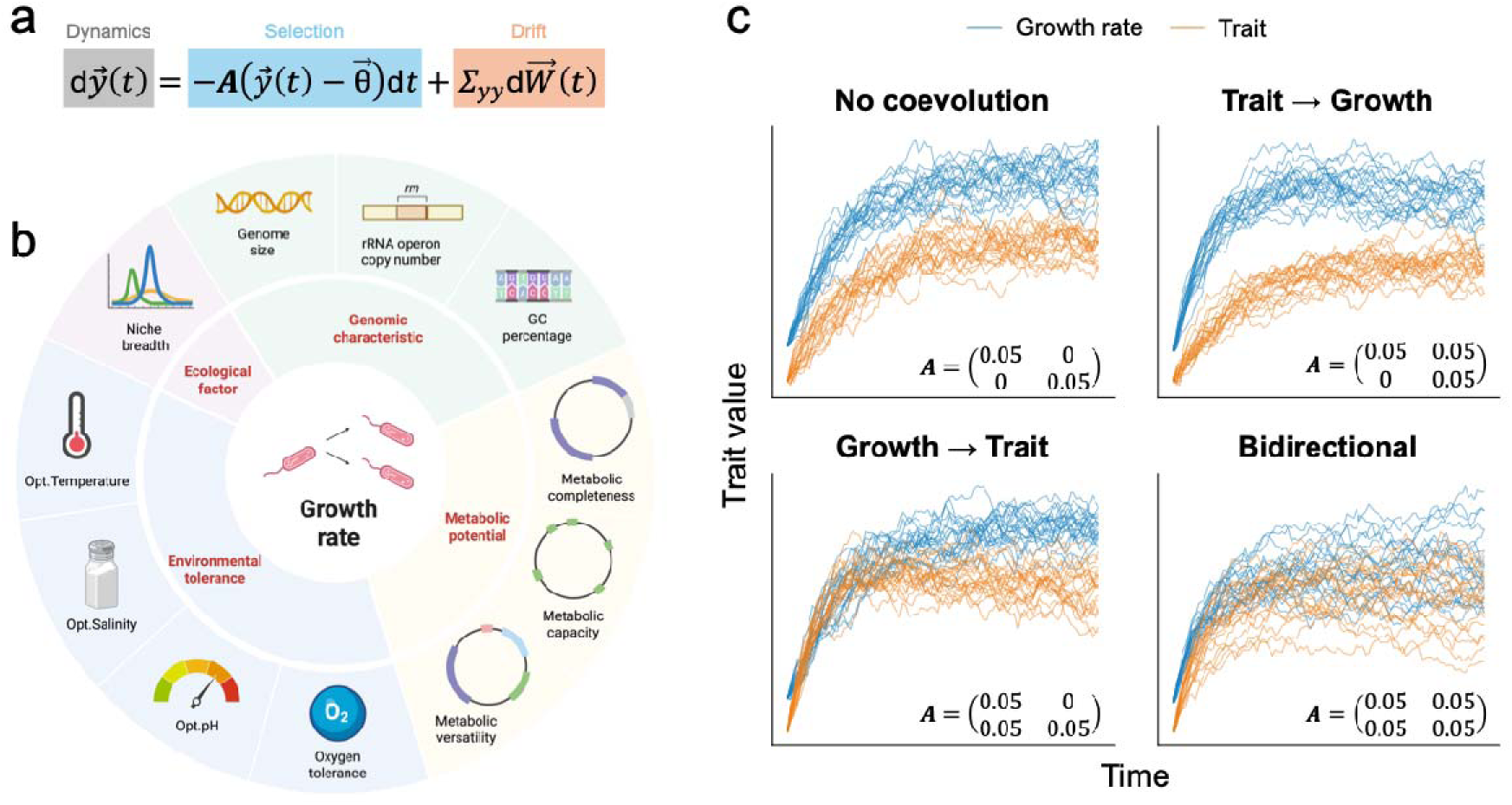
Conceptual framework and hypotheses. **(a)** The multivariate Ornstein–Uhlenbeck (mvOU) process models joint trait evolution as deterministic attraction of the trait vector 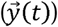 towards adaptive optima 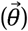, subject to stochastic drift 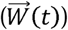. The diagonal elements of adaption matrix *A* quantify self-adaptation of each trait towards its optimum, while off-diagonal elements quantify how the evolution of one trait is influenced by the state of another trait, determining the strength and direction of their coevolution. Details of the model framework is given in Methods. **(b)** Eleven genomic, metabolic, ecological and environmental traits hypothesised to shape the evolution of bacterial growth rate. Details of these hypotheses and current empirical support for the effect of each trait were summarized in Table S1. **(c)** Simulated evolutionary trajectories under four alternative coevolutionary modes between growth rate (blue) and an influencing trait (orange). Simulations were run under a bivariate OU process with 20 replicate trajectories run from a shared ancestral state set-up (*y_0_* = (−1.0, −1.5) towards fixed optima (*θ* = (1.5,0.5), over 100-time units. Coevolutionary mode was determined by the structure of *A*, with diagonal elements fixed at 0.05 and off-diagonal elements set to 0 or 0.05 to represent the absence, direction or reciprocity of coupling. Stochastic variation was modelled with diagonal noise covariance, 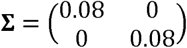. These scenarios define the alternative evolutionary hypotheses tested in this study.

## Results

### Data overview

We compiled experimentally measured minimal doubling times for 283 bacterial species from the Madin et al. trait database^38^ (Table S6). The distribution of doubling times was highly right-skewed, indicating that most species grow rapidly while a few exhibits very slow growth (Figure 2a). To aid interpretation, we expressed maximum growth rate (hereafter “growth rate”) as the negative logarithm of the minimal doubling time, such that higher growth rates correspond to shorter doubling times (Figure 2b). Growth rate in our dataset showed a moderate but significant phylogenetic signal, with Blomberg’s *K* statistic^39^ = 0.205 (*P* < 0.001, 1,000 randomizations) and Pagel’s *λ* statistic^40^ = 0.694 (*P* < 0.001, likelihood-ratio test; Figure 2c & Table 1).

For each species, we also assembled data on 11 traits proposed to affect bacterial growth rate (Figure 1b; Table S6). These included: three genomic traits — genome size (kb), GC percentage (%), and ribosomal RNA operon (*rrn*) copy number; three metabolic traits — metabolic capacity, metabolic versatility, and metabolic completeness (see Methods for definitions); four traits related to environmental tolerance — optimum temperature (°C), optimum salinity (% w/v NaCl), optimum pH, and oxygen tolerance (probability of tolerance ranging from 0 for strict anaerobes to 1 for strict aerobes); and one ecological trait — niche breadth, defined as the diversity of habitats occupied by each species (Figure 1b & 2c; Table S1). Trait values were standardized (mean = 0, SD = 1) to allow direct comparison among traits with different units and ranges (Figure 2c; Table S7). We found that all of these traits exhibited significant phylogenetic signal, whereas growth rate displayed the weakest phylogenetic signal among all measured traits (Table 1; Figure 2c).

**Figure 2.**
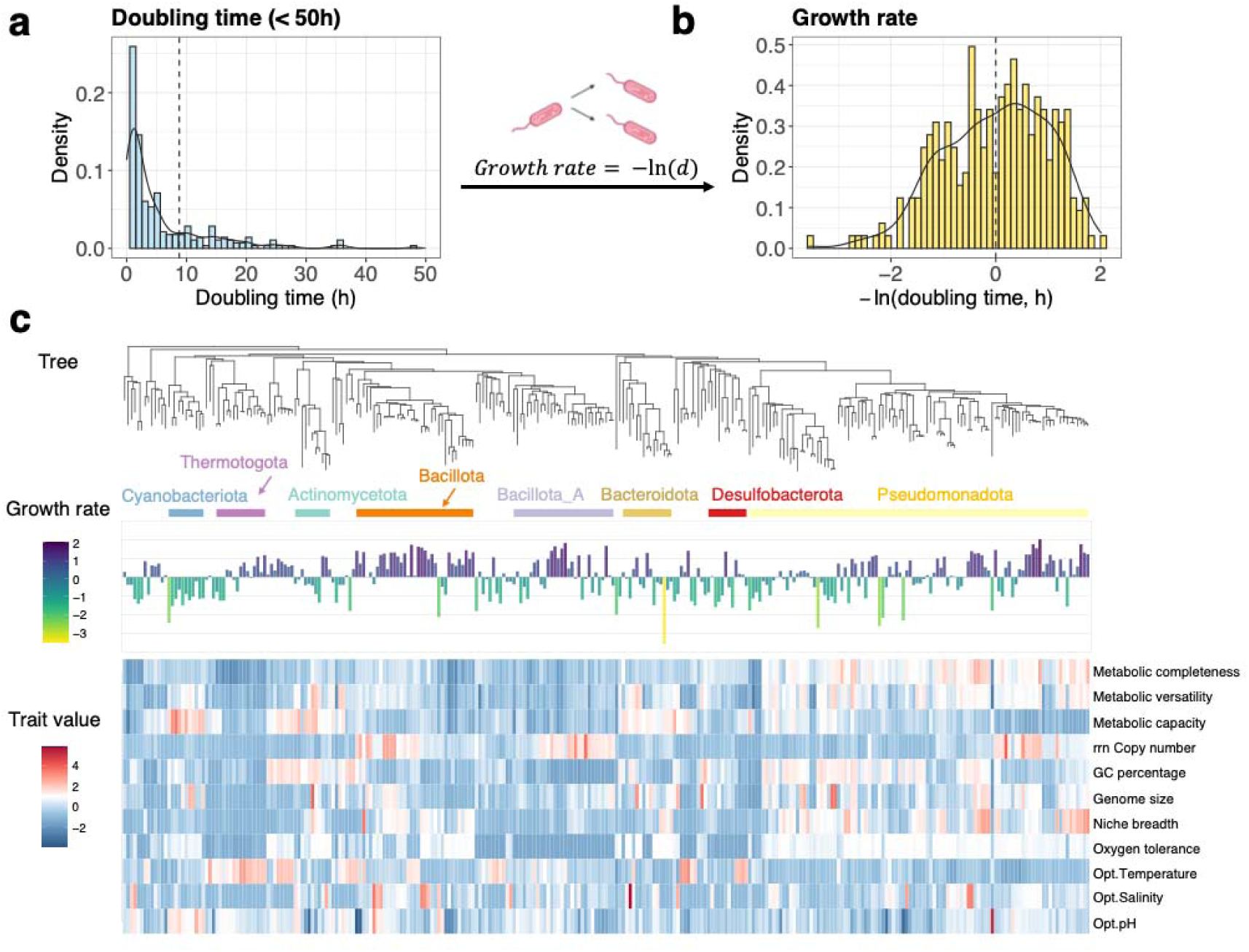
Data overview. **(a)** Distribution of doubling times for 283 bacterial species with experimentally measured growth rates (values truncated at 50 h for visualisation). **(b)** Corresponding distribution of growth rate, expressed as *ln(doubling time)*, used throughout the study. **(c)** Bacterial phylogeny annotated with phylum-level taxonomy (top), growth rates mapped as bar plots (middle; bar height and colour intensity proportional to growth rate), and a heatmap of the eleven standardised traits (bottom), illustrating the phylogenetic diversity represented in the dataset.

### Growth rate evolution is best described by an Ornstein–Uhlenbeck process

We first modelled the evolution of individual traits. We compared alternative models of trait evolution, including Brownian motion (BM), Ornstein–Uhlenbeck (OU), and early burst (EB), to characterize the mode of evolution for growth rate, and found that the OU model provided the best fit (lowest AICc; Figure S1). This pattern is consistent with growth rate evolution being constrained around an adaptive optimum, rather than by neutral drift (BM) or adaptive radiation (EB)^30,34^.

When modelling the evolution of the 11 associated traits, we found that 7 were best described by the OU model, 3 by the EB model, and 1 by the BM model, suggesting that most traits evolved under stabilizing evolutionary processes, whereas a few exhibited time-dependent or neutral evolutionary dynamics (Table 1; Figure S1). For the traits whose evolutionary mode was best described by an OU process, we compared their evolutionary half-lives (*t*_1/2_) — defined as the time required for the expected trait value to move halfway from its ancestral value toward the optimum, to compare the rate of evolution across these traits^29^. Growth rate had the shortest half-life among these traits, corresponding to 25.5% of the total tree height, suggesting that it returns to its optimum most rapidly (Table 1). This is consistent with its comparatively weak phylogenetic signal, as rapid attraction toward the optimum reduces the persistence of phylogenetic similarity (Table 1).

To further evaluate the robustness of these evolutionary mode and tempo estimates, we independently repeated the analysis using bioinformatically estimated trait values from a larger dataset encompassing 24,402 species (Figure S12). The inferred evolutionary modes and OU half-life parameters showed consistent patterns between the two datasets, supporting the robustness of our findings (Figures S14 & S15).

**Table 1.**
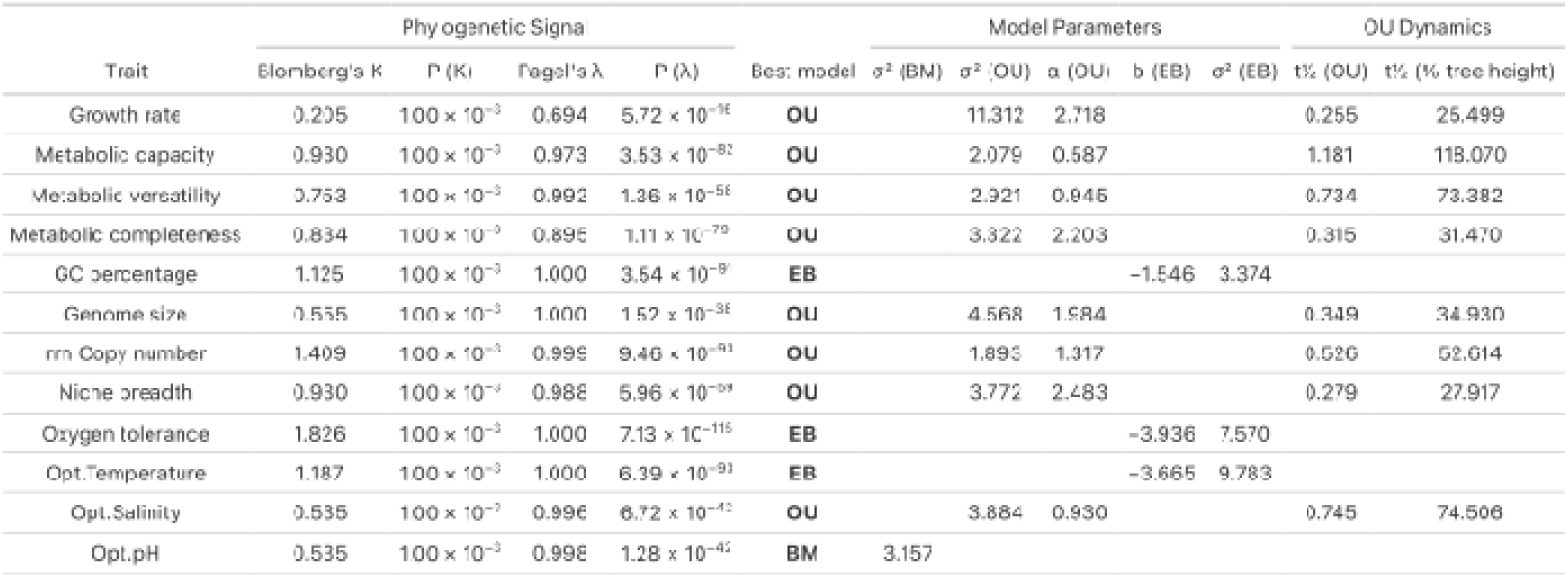
Phylogenetic signal and trait-evolution model fitting for each individual trait. Two metrics were computed to quantify phylogenetic signal — Blomberg’s *K* and Pagel’s λ. For both metrics, statistical significance was evaluated using the randomization test. A p-value ≤ 0.05 indicates that the observed trait exhibits a phylogenetic signal stronger than expected by chance. All traits analysed here show significant phylogenetic signal. The best evolutionary model for each trait is identified as either BM, OU, or EB, with the corresponding model parameters displayed for each model. Parameter interpretations — σ^2^(BM), σ^2^(OU), σ^2^(EB): the variance of the evolutionary process for each model, describing how much the trait is expected to change over time; α(OU): the rate of adaption of a trait under the OU process; *b*(EB): the decay parameter of EB model, determining how quickly the rate of evolution decreases over time; *t*_1/2_(OU): The half-life (*t*_1/2_) of OU models, representing the time required for a trait to evolve halfway to its optimal state. The *t*_1/2_ is calculated using the formula: 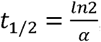, and expressed as a fraction of total tree height. Growth rate turns out to have the smallest *t*_1/2_.

### Multiple selective regimes underlie the evolution of growth rate and several traits

We subsequently used a Bayesian OU modelling framework — *bayou*, to test whether the OU dynamics of these traits inferred above are better described by evolution toward a single global adaptive optimum (*θ*; OU1) or by shifts among multiple optima^41^ (OUM; see in Methods). For growth rate, Bayes factor comparisons based on stepping-stone marginal likelihoods strongly favoured the OUM model (*logBF_OUM-OU1_* = 7.55; Figure 3a; Table S2). Across MCMC chains, the mean estimated number of selective regimes was 2.04 (Table S2), suggesting approximately two growth-rate regimes across the phylogeny. We therefore identified the node with the highest posterior probability of a regime shift and used it to define two distinct selective regimes for growth rate: a slow-growing regime (*θ* = −0.157), and a fast-growing regime (*θ* = 0.582; Figure 3b; Figure S2; Table S3). Examination of the taxonomic distribution of regimes showed that the fast-growing regime was primarily associated with the phylum *Bacillota* (Figure 3b).

Extending this analysis to the 11 associated traits revealed trait-specific support for regime shifts. OUM was favoured for 6 traits, whereas OU1 was favoured for 5 traits (Figure 3a; Table S2). For the 7 traits (including growth rate) favouring OUM, models inferred only a few distinct optima (posterior means ≈ 2.0 to 2.6), with 95% highest posterior density (HPD) intervals typically spanning 1 to 4 optima, indicating that the supported multi-optimum signal reflects a limited number of recurrent regime shifts rather than many regimes (Figure 3a; Figure S3; Table S2). Because growth rate and several traits show strong evidence for regime shifts when modelled independently, we next test whether incorporating regime-dependent dynamics improves inference of the coevolutionary relationships between growth rate and its associated traits.

**Figure 3.**
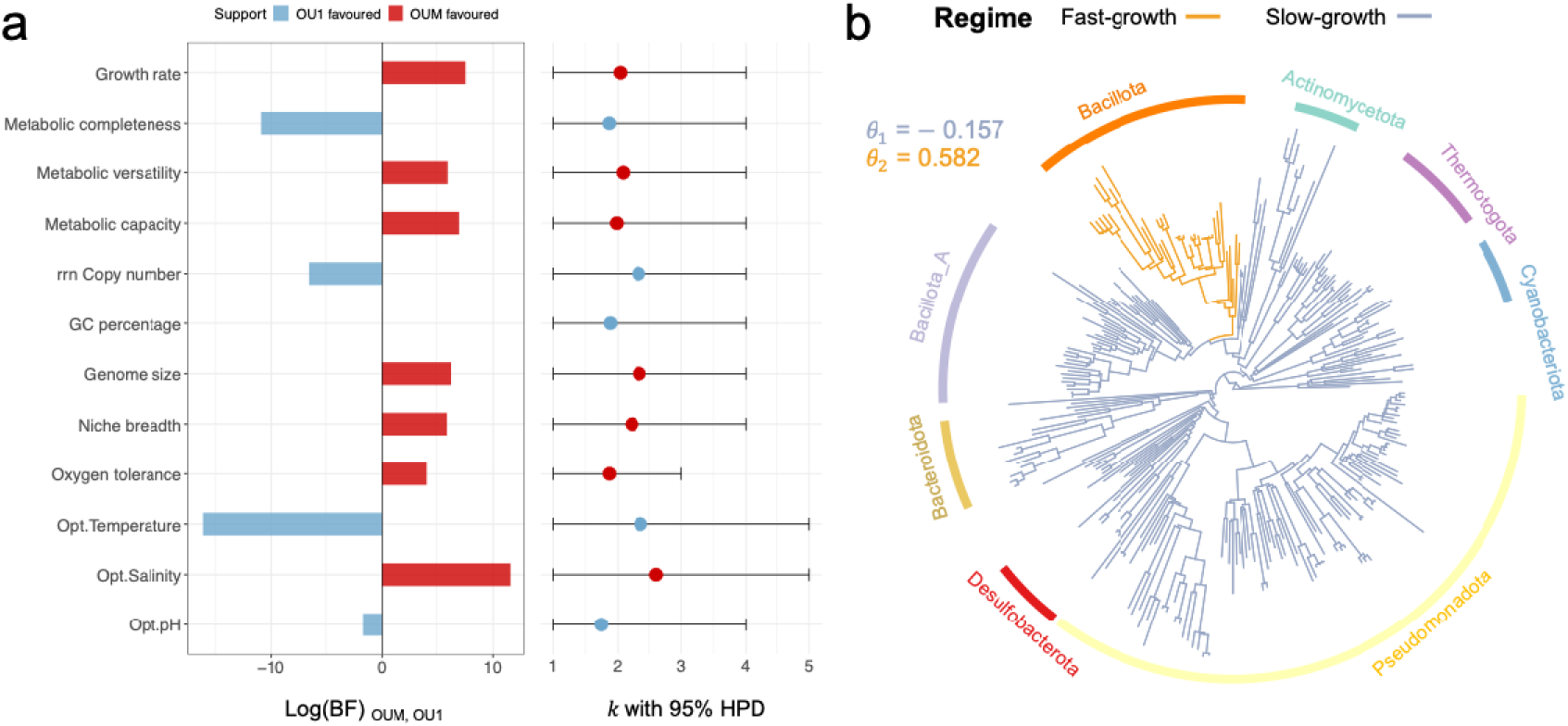
Results of *bayou* reversible-jump MCMC analyses under a multi-regime OU framework. **(a)** Comparison of single-(OU1) and multi-optimum (OUM) Ornstein–Uhlenbeck models across traits. For each trait, evolutionary model support was evaluated by comparing a OU1 model with a OUM model using Bayes factors. Left panel: horizontal bars show the log Bayes factor, calculated as the difference between the log marginal likelihoods of the OUM and OU1 models. Positive values (red bars) indicate support for the OUM model, whereas negative values (blue bars) indicate support for the OU1 model. Right panel: points and horizontal lines show posterior means and 95% highest posterior density (HPD) intervals for number of selective regimes (*k*) estimated under the OUM model. **(b)** Phylogenetic distribution of selective regimes for growth rate. The slow-growing regime corresponds to the root regime, whereas the fast-growing regime is primarily associated with the phylum *Bacillota*.

### Coevolution of growth rate and traits is primarily structured by growth rate regimes

Based on the results of the univariate model fitting, we next investigated the mode of coevolution between growth rate and each of the 11 associated traits using multivariate trait evolution models. For each trait pair, we first modelled their coevolution under a single-regime OU process (OU1), in which both traits evolve toward constant, trait-specific optima, and compared model fit to a corresponding Brownian motion model (BM1) that assumes neutral evolution for both traits. In all cases, the OU1 models provided a consistently better fit (lower AICc) than the BM1 models, indicating that the coevolution of growth rate with these traits is better explained by stabilizing evolution rather than neutral drift over macroevolutionary time (Figure S4).

We next examined the coevolution of each trait pair by comparing the multi-regime (optima) OUM models to the single-regime OU1 models under the multivariate Ornstein–Uhlenbeck (mvOU) framework^30,32,42^. For each pair, we evaluated three regime settings: (i) the growth rate regime, (ii) the regime of the associated trait, and (iii) the combined regime of both growth rate and the trait. To ensure robustness, we modelled different levels of measurement error and reported the best-supported pattern across different measurement-error settings for the coevolution of growth rate with each trait^43,44^ (see Methods). We found that, when stochastic perturbations between the two traits were assumed to be independent (i.e., when the *Σ_yy_* matrix was constrained to be “Diagonal”; see Methods), OUM models provided a better fit than OU1 models across all trait pairs and measurement-error settings. This indicated that the coevolution of growth rate and its associated traits is better explained by distinct adaptive regimes than by a single shared optimum (Figure 4; Figure S5; Table S4).

When comparing the best-supported models across measurement-error settings for each trait pair (Figure S6), the coevolution of growth rate and the associated trait is primarily driven by the growth rate regime, except for niche breadth, where the coevolution is shaped by the combined regimes of both growth rate and niche breadth (Figure 4; Table S5). Moreover, the finding that growth-rate regimes primarily structure these coevolutionary dynamics held when allowing correlated stochastic perturbations between the two traits^42^ (i.e., “UpperTri” *Σ_yy_* matrix; Figure S7). We also observed no substantial difference in model fit between the two *Σ_yy_* matrix assumptions (Figure S8). We therefore focused on results obtained under the “Diagonal” assumption, which avoids unnecessary model complexity without compromising the robustness of the conclusions.

**Figure 4.**
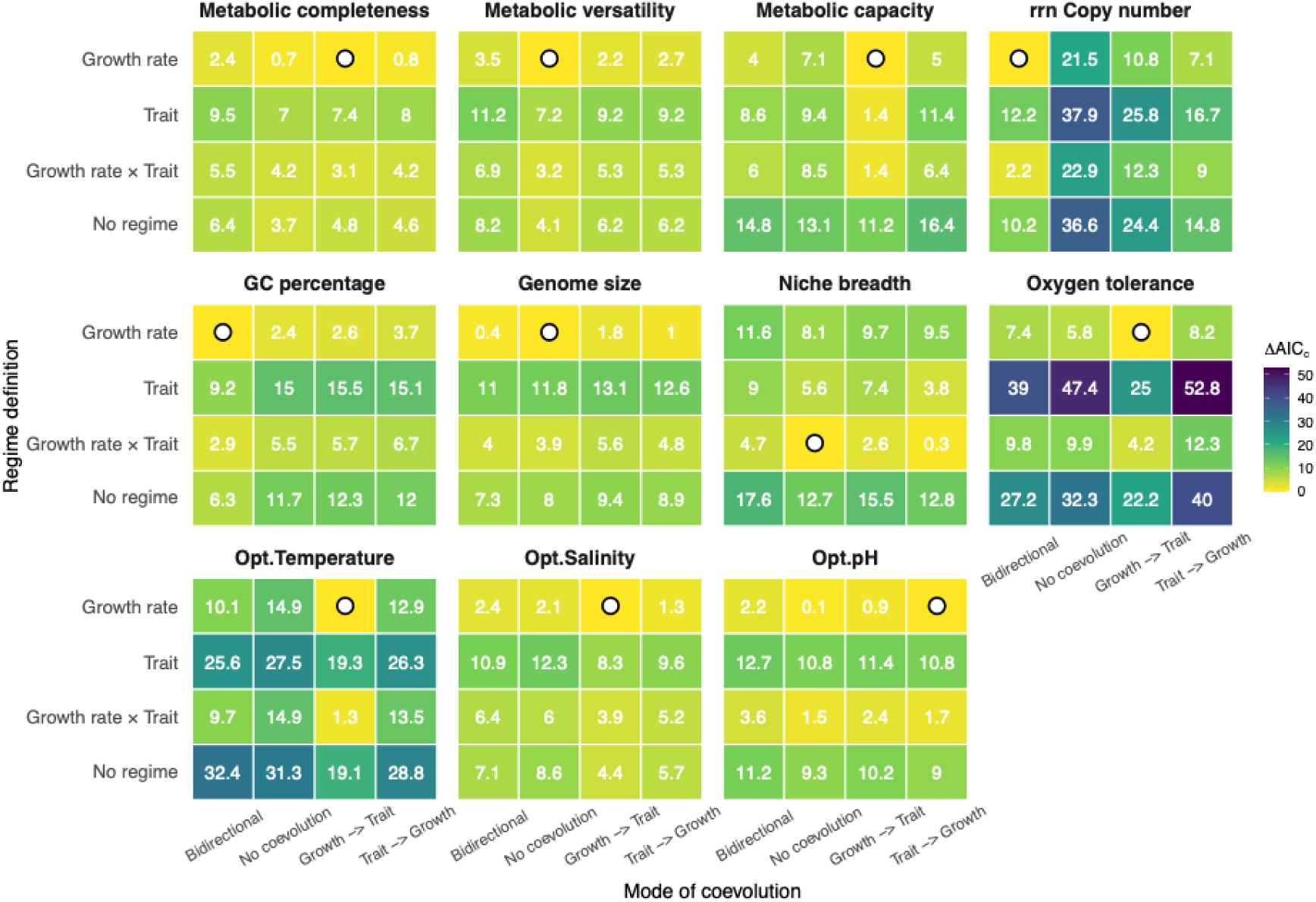
Mode of coevolution between growth rate and each trait. Each panel represents the comparison of goodness-of-fit for bivariate OU models between growth rate and one of the 11 associated traits under different evolutionary modes. The x-axis shows the presence/absence and directionality of coevolution (i.e., mode of evolution), while the y-axis represents the regime definitions under which coevolution is evaluated. For each combination of mode and regime, the best-supported measurement-error setting (lowest AICc) was retained (see Figure S6 for the comparison of goodness-of-fit across measurement error levels). The heatmap colours represent the ΔAICc values, with lower values (closer to zero) indicating better model fits. The circle inside a grid indicates the model with the best fit (ΔAICc = 0) for the coevolution between growth rate and that trait pair. Results are shown for models in which the *Σ_yy_* matrix was constrained to a “*Diagonal*” structure, thereby assuming independent stochastic perturbations between the two traits.

### Growth rate coevolves with associated traits through distinct tempos and modes

To further characterize the mode of coevolution between growth rate and each trait over instantaneous timescales, we compared mvOU models with different adaptation matrix (*A* matrix) structures, distinguishing among independent evolution (no coevolution), unidirectional coupling, and bidirectional co-adaptation (Figure 1c; Figure S9). Based on the best-supported models across measurement-error settings for each trait pair (Figure 4), the evolution of growth rate with metabolic versatility, genome size, and niche breadth was best explained by uncoupled evolution. In contrast, growth rate exhibited bidirectional coevolution with *rrn* copy number and GC percentage. For the remaining traits, unidirectional models were supported, with asymmetry in the direction of evolutionary influence. Specifically, the best-supported models indicated directional evolutionary dependence from optimum pH to growth rate (trait → growth rate), whereas for the other five traits, the supported direction was from growth rate to the associated trait (growth rate → trait; Figure 5a; Table S5).

Examining the tempo of coevolution, we first investigated the rate of adaptation over instantaneous evolutionary timescales, as reflected by the estimated *A* matrix coefficients (Figure S9; see Methods). Our analysis revealed that similar modes of coevolution can correspond to different instantaneous rates of co-adaptation. For instance, for *rrn* copy number, deviations from its optimum constrained growth rate (rate = −13.88), whereas deviations in growth rate promoted adaptive shifts in *rrn* copy number toward its optimum (rate = 5.10; Figure 5a). In contrast, for GC percentage, deviations in GC content drove growth rate toward its optimum (rate = 8.33), while deviations in growth rate constrained GC percentage away from its optimum (rate = − 6.19; Figure 5a).

By eigen-decomposing the fitted *A* matrices, we examined how linear combinations of growth rate and each trait evolve along independent adaptive axes over extended evolutionary timescales^45,46^. Each eigenvector defines an axis of co-adaptation of two traits in the joint trait space. The corresponding eigenvalue quantifies the strength of adaptive pull along that axis over longer timescales, with larger values indicating faster stabilization toward the optimum (Figure S9; see Methods). We summarized these tempos as evolutionary half-lives, expressed relative to total tree height (Figure 5b; Figure S10).

Across all models, growth rate consistently dominated the fast evolutionary mode 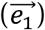, indicating relative rapid evolution toward its adaptive optimum over longer evolutionary timescales. In contrast, the associated traits often loaded more heavily on the slow mode 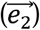, reflecting more gradual, inertia-like adaptive responses (Figure 5b; Figure S10). This pattern was consistently observed when modelling the coevolution of growth rate with multiple composite trait axes, allowing us to test whether growth-rate evolution is influenced by the combined effects of genomic, metabolic, ecological, and environmental traits (Figure S26).

Moreover, for metabolic versatility, genome size and niche breadth, 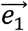 and 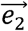 were orthogonal (i.e., growth rate completely dominates 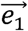 whereas the associated trait completely dominates 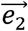), indicating that these traits evolved independently of growth rate across deep evolutionary time (Figure 5b; Figure S10).

**Figure 5.**
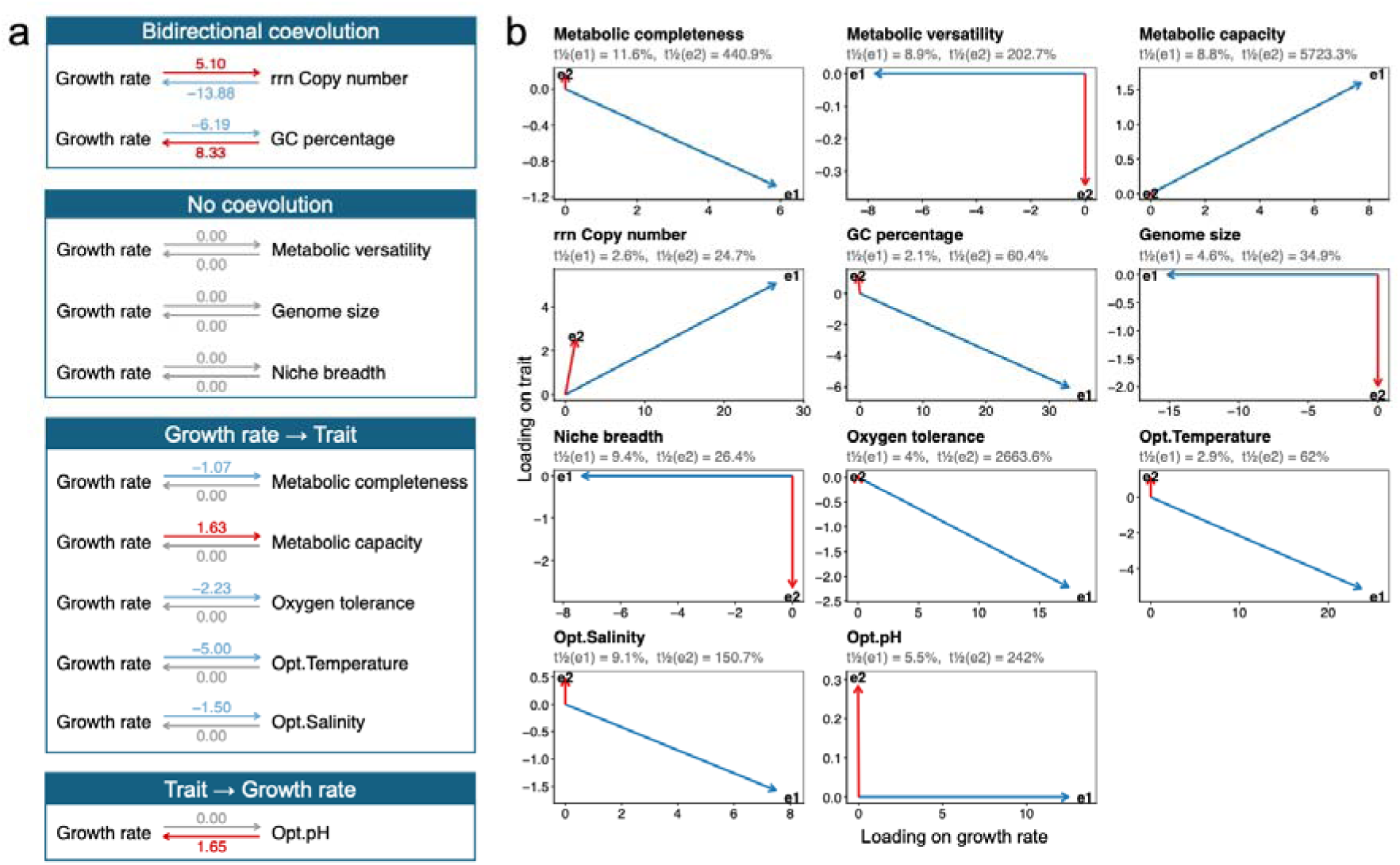
Tempo and modes of coevolution at distinct temporal scales. **(a)** The off-diagonal structure of the *A* matrix for the best-supported model of each trait pair, distinguishing independent evolution, unidirectional coupling, and bidirectional co-adaptation. Arrows denote the direction and strength of adaptive coupling, where a change in one trait induces an evolutionary response in another. Red arrows represent adaptive pull toward an optimum, whereas blue arrows indicate deviation away from the optimum. Coefficients on arrows indicate the rate of adaptation at instantaneous evolutionary timescales. **(b)** Eigen-decomposition of the *A* matrix, illustrating evolutionary dynamics over longer timescales. Arrows represent the two eigenvectors of the *A* matrix 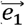 in blue, 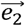 in red), plotted in trait space defined by growth rate (x-axis) and the focal trait (y-axis). The direction of each arrow corresponds to the loadings of the eigenvector on growth rate (horizontal component) and the focal trait (vertical component), defining the combination of traits that evolve together along that evolutionary mode. Positive and negative loadings indicate whether traits change in the same or opposite directions along the eigenvector, with the overall sign defining the orientation of the evolutionary axis in trait space. Arrow length is scaled by the corresponding eigenvalue (*λ*), such that longer arrows represent faster evolutionary modes, whereas shorter arrows represent slower modes. Accordingly, 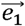 generally corresponds to the fastest evolutionary axis (largest λ), and 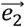 to the slowest. The tempos of each mode were summarized as evolutionary half-lives 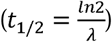 expressed as a fraction of total tree height. For detailed values, refer to Figure S10.

## Discussion

Using a multivariate OU framework, we revealed that the coevolution of bacterial growth rate with its associated traits exhibits both a consistent macroevolutionary asymmetry—growth rate generally evolves more rapidly and its regimes primarily structure coevolution—and substantial trait-specific variation in evolutionary coupling, patterns not readily captured by conventional comparative methods.

Traditional methods such as PGLS are useful for detecting whether traits covary across extant species, but they do not identify the macroevolutionary processes that generate those covariation patterns^47^. Our simulation illustrates this limitation: different macroevolutionary dynamics, ranging from unidirectional influence to bidirectional coevolution or independent evolution, can produce similar apparent correlations among extant species (Figure S17). Consistently, the mvOU models recovered non-zero stationary covariance for eight of the eleven trait pairs, compared with only three significant associations under PGLS, showing that traits with little detectable covariation among extant species may nonetheless remain evolutionarily coupled through processes that generate long-term covariance (Figures S16 and S19; see Supplementary analysis 2).

Our bivariate mvOU analyses showed that the coevolution of growth rate with associated traits is primarily structured by growth-rate regimes (Figure 4). This pattern reflects fundamental trade-offs in cellular resource allocation that constrain how other traits can evolve^10,48^. Studies on microbial proteome economics show that cells face trade-offs between maximizing growth and investing in adaptability, stress resistance, or resource uptake^11,49–52^. For example, nutrient-rich conditions favour allocation toward ribosome synthesis and central metabolism to maximize growth, while nutrient limitation favours investment in high-affinity uptake and stress-response systems at the expense of rapid growth^48,51^. These contrasting strategies underpin the classic copiotrophic versus oligotrophic dichotomy, which may shape how other microbial life-history strategies evolve over macroevolutionary time^53,54^. Growth-rate regimes may therefore capture broader differences in cellular and ecological strategy that jointly structure the evolutionary trajectories of multiple associated traits. Such regime dependence also illustrates why a single cross-species association can be incomplete: a single PGLS slope across lineages from different adaptive regimes conflates distinct regime-specific coevolutionary relationships and directional asymmetries that mvOU models explicitly resolve.

Niche breadth is an exception in our analyses: unlike most traits, its coevolution with growth rate was best explained by a combined regime incorporating both growth-rate and niche-breadth optima (Figure 4). This likely reflects the integrative, emergent nature of niche: niche is not a single physiological or genomic trait but an outcome of multiple interacting constraints, including metabolism, environmental tolerance, and habitat specificity^55–59^. Thereby, niche breadth defines the ecological context in which growth can occur. By determining the range of environments a lineage can exploit, niche breadth constrains which growth-rate strategies are viable. Consequently, the adaptive optimum of growth rate depends on both the lineage’s intrinsic growth strategy and the ecological opportunities imposed by niche breadth, producing the combined regime observed in our analyses.

We also revealed that different traits coevolve with growth rate through distinct modes. For example, although genome size is often proposed as a major constraint on bacterial growth^20–22^, we found that the macroevolutionary trajectories of genome size are largely independent of growth rate at both instantaneous and extended timescales (Figure 5). One possible explanation is that genome size evolution in bacteria can be driven by a variety of events, such as gene duplication, gene deletion, and horizontal gene transfer^60–63^. These evolutionary events are often episodic and operate at varying tempos rather than continuously^64^, potentially decoupling genome size from the more continuous selective pressures influencing growth rate evolution.

In contrast, *rrn* copy number showed a more dynamic and reciprocal relationship with growth rate (Figure 5). Prior studies using PGLS have noted a strong positive correlation between growth rate and *rrn* copy number across species^65^, which is also recovered by our additional analysis (Figure S16). But our analysis goes further by uncovering the underlying evolutionary processes. We found a feedback loop where *rrn* copy number not only responds to but also drives changes in growth rate, a pattern that further reinforces the interdependence of these traits (Figure 5a). In addition, we further revealed that both traits evolve in a coordinated manner at longer timescales, moving in the same direction. This suggests that the coevolution between these two traits becomes even more tightly integrated over macroevolutionary periods, underscoring the importance of reciprocal interactions in shaping trait evolution (Figure 5b).

By showing that growth rate evolves fastest (i.e., having the shortest evolutionary half-time or dominating the fastest-evolved evolutionary modes) over extended macroevolutionary timescales across univariate (Table 1), bivariate (Figure 5b), and multivariate (Figure S26) OU models, our analyses suggested that growth rate functions as a macroevolutionary pacemaker: it shifts rapidly toward adaptive optima, while other traits display more inertia-like evolution around those shifts. This is likely because growth rate directly influences organismal fitness and ecological success, which makes it highly responsive to environmental pressures^1,3,4^.

Some limitations and interpretation caveats should be considered. First, mvOU models infer evolutionary coupling from comparative data; they do not directly demonstrate mechanistic causation^31,32,42,66,67^. The directionality of off-diagonal terms should therefore be interpreted as evidence for asymmetric evolutionary dependency rather than direct experimental cause and effect. Second, experimentally measured minimal doubling times are compiled from heterogeneous conditions and may not always reflect the realized growth rates of species in natural environments^15,38^. Although our modelling framework incorporated alternative measurement-error settings to partly account for uncertainty in growth-rate estimates, this approach cannot fully resolve variation arising from differences in experimental protocols, culture conditions, or ecological contexts. Third, the dataset necessarily emphasizes cultured and experimentally characterized taxa, which may underrepresent slow-growing, uncultured or environmentally specialized bacterial lineages^15,13,38^. Where applicable, we assessed the robustness of our findings using a much larger dataset based on bioinformatically predicted growth rates, thereby partially mitigating potential biases associated with limited taxonomic coverage (Figure S12). Fourth, species-level trait values can mask substantial strain-level variation^68–71^. To address these limitations, future work should focus on integrating growth rate data with richer ecological metadata, including environmental abundance data and more granular strain-level genome variation. Additionally, standardized growth measurements under experimentally controlled conditions would allow for better comparison across species and provide more robust insights into the observed macroevolutionary patterns.

## Methods

### Growth rate data compilation

We compiled experimentally measured minimal doubling times (in hours) for bacterial species from the Madin et al. trait database^38^. Species whose scientific names did not match those of the compiled GTDB species (see below) were excluded. To facilitate interpretation, we defined the maximum growth rate (hereafter referred to as growth rate) as the negative logarithm of the minimal doubling time for each species, such that species with higher growth rates have shorter doubling times (Figure 2b).

Bioinformatically estimated maximum growth rates could, in principle, expand taxonomic coverage. However, when we compared growth-rate estimates from two commonly used resources, Phydon^13^ and gRodon^15^, against experimentally measured minimal doubling times, we found limited agreement between predicted and measured values (Figure S11). Because multivariate OU models can be sensitive to measurement error, we restricted our main analyses to species with experimentally measured growth-rate data^43,44,72^. This conservative approach reduced the number of species included in the analysis but minimized the risk that prediction error in growth rate would bias inference of evolutionary tempo, adaptive optima, or trait coupling. Bioinformatically estimated growth rates were therefore used only as a robustness check for exploring evolutionary modes across traits in this study (Figure S13-15).

### Genome collection

We obtained representative genomes of each of the species in our dataset from the Genome Taxonomy Database (GTDB^73^; release 220) on 1^st^ July 2024. For these species, we adopted a stricter standard than the Minimum Information about a Metagenome-Assembled Genome (MIMAG) criterion^74^, retaining only high-quality representative genomes with completeness ≥ 95% and contamination ≤ 5%. Nucleotide and amino acid sequences for all representative genomes were downloaded using *ncbi-genome-download* scripts (version 0.3.1) from either GenBank^75^ or RefSeq^76^, based on GTDB assembly entries.

### Trait definition and data processing

We identified 11 traits that are proposed to influence bacterial growth rates in various ways, these traits represent genomic, metabolic, ecological, and environmental constraints on growth (Figure 1b; Table S1). Empirical evidence for the relationship between each trait and growth rate is summarized in supplementary materials, showing that the expected associations vary across studies and can be weak, absent, or even reversed (Table S1). We compiled data for all traits across the previously assembled species list, yielding a final dataset of 283 species for analysis (Table S6).

#### (1) Genomic characteristic

We identified three genomic characteristics that have been proposed to influence bacterial growth rate: genome size, GC percentage, and ribosomal RNA operon (*rrn*) copy number (Figure 1b; Table S1). Genome size (kb) and GC percentage (%) for each species were calculated directly from their representative genomes. RNA operon copy number data were obtained from the *rrnDB*^77^, with missing values estimated by phylogenetic imputation from related taxa.

#### (2) Metabolic potential

We identified three traits representing the metabolic potential of bacterial species: metabolic capacity, metabolic versatility, and metabolic completeness (Figure 1b; Table S1). Metabolic capacity was defined as the proportion of metabolic genes encoded in a species’ representative genome, reflecting its overall ability to perform metabolic reactions and processes. Metabolic versatility was defined as the number of distinct metabolic pathways encoded by the representative genome, indicating the potential to utilize a wide range of compounds as energy or nutrient sources^78^. Metabolic completeness (%) was defined as the average completeness of metabolic pathways encoded in the representative genome, reflecting both a species’ potential for autonomous biosynthesis and its tendency toward auxotroph^79,80^.

The metabolic genes and pathways were identified using the KEGG Orthology (KO) database^81^. KO identifiers were assigned to protein sequences using KofamScan^82^. We defined metabolic KOs as those mapped to pathways within the “Metabolism” category of KEGG PATHWAY Database^81^, which includes 189 pathways grouped into 13 different classes. Associations between KO (prefixed “ko_”) to pathway (prefixed “map_”) were retrieved via KEGG API and used to assign each identified KO to its corresponding pathway across all genomes (https://rest.kegg.jp/link/ko/pathway). Genes that could be assigned to at least one metabolic KO were considered as metabolic genes, and their linked pathways were considered metabolic pathways. We used the MGnify KEGG Pathways Completeness Tool^83^ to calculate the completeness of each KEGG pathway module (n = 495) based on the presence or absence of the corresponding KOs in each genome. For each species, metabolic completeness was computed as the average completeness of all KEGG pathway modules present in its representative genome.

#### (3) Environmental tolerance

We considered four traits reflecting environmental constraints on bacterial growth rate (Figure 1b; Table S1): optimum temperature (°C), optimum salinity (% w/v NaCl), optimum pH, and oxygen tolerance (probability of tolerance, ranging from 0 for strict anaerobes to 1 for strict aerobes). Environmental tolerance data for each species were compiled from the GenomeSPOT database (https://github.com/cultivarium/GenomeSPOT), which uses a genome-based computational model to predict microbial growth conditions from amino acid composition^84^.

#### (4) Niche breadth

Niche breadth has also been proposed to influence bacterial growth rates (Figure 1b; Table S1). In its strict definition, niche breadth describes the range of biotic and abiotic conditions under which a species can maintain positive fitness^56,85^. However, such direct quantification is rarely feasible, and microbial ecology studies typically rely on proxies. Here, we defined the niche breadth of a bacterial species as the diversity of habitats (e.g., water, soil, etc.) it can inhabit, consistent with common practice in microbial ecology^59,86^.

To estimate habitat preferences for each species, we retrieved full-length 16S rRNA gene sequences from their representative genomes in GTDB. These sequences were queried using BLAST against 16S rRNA sequences from metagenomic datasets with associated environmental metadata through the ProkAtlas online platform^87^. Sequence similarity and coverage thresholds were set to 97% and 150 bp, respectively, for ProkAtlas annotation. A species was considered present in an environment if its habitat preference score reported by ProkAtlas was greater than zero (Table S8). Niche breadth was then calculated as the number of habitats in which a species was detected.

#### (5) Data standardization

To ensure comparability among traits with different units and ranges, all traits were standardized prior to multivariate analyses to have a mean of 0 and a standard deviation of 1 (Figure 2c; Table S7). This data is used for our main analyses.

### Phylogeny

The reference phylogenetic tree of bacterial species was obtained from GTDB^73^ (release 220). We pruned this tree to include only the species in our dataset using the “*keep.tip*” function in the R package *ape* (version 5.7.1)^88^, resulting in the phylogenetic trees used in subsequent analyses (Figure 2c).

### Phylogenetic signal

We first evaluated the phylogenetic signal of bacterial growth rate and each of the 11 associated traits. A strong phylogenetic signal indicates that trait values are evolutionarily conserved, such that closely related species resemble each other more than expected by chance^39^. We quantified phylogenetic signal using Pagel’s λ^40^ and Blomberg’s *K*^39^ for experimentally measured growth rates and associated traits, implemented with the “*phylosig*” function in the R package “*phytools*”^89^ (version 2.5.2). For both metrics, statistical significance was evaluated using the built-in randomization test. Specifically, the observed value of *K* or *λ* was compared against a null distribution generated by randomly reshuffling the trait values across the tips of the phylogeny. The p-value represents the proportion of randomizations that produce a statistic equal to or more extreme than the observed value. A p-value ≤ 0.05 indicates that the observed trait exhibits a phylogenetic signal stronger than expected by chance.

### Characterizing the mode of trait evolution

To characterise the mode of evolution for growth rate and each associated trait, we compared the fit of three standard models of continuous trait evolution: Brownian motion (BM), Ornstein–Uhlenbeck (OU) and early burst (EB)^47^. BM models evolution as a random walk with variance increasing linearly through time, reflecting unconstrained drift. OU models incorporate stabilising evolution toward an adaptive optimum, characterised by an attraction parameter (*α*) that quantifies the strength of the pull toward the optimum (*θ*). EB models allow evolutionary rates to decrease or increase exponentially through time, capturing patterns such as early rapid diversification followed by slowdown^47^. We fitted each model using the *“fitContinuous”* function in the “geiger” R package^90^ (version 2.0.11), with 100 random starting points employed to ensure more precise likelihood estimation. Model support was assessed using the corrected Akaike information criterion (AICc), with lower AICc values indicating better fit. For the best-supported model of each trait, we extracted the relevant model parameters. σ^2^_BM_, σ^2^_OU_, and σ^2^_EB_ represent the variance in the evolutionary process for each model, describing how much the trait is expected to change over time. For the EB model, parameter *b* represents the decay parameter, which determines how rapidly the rate of evolution decreases over time. For the OU model, *α* is the rate of adaption of a trait under the OU process. We also calculated the half-life (*t*_1/2_) of OU models, representing the time required for a trait to evolve halfway to its optimal state^29,30^. The *t*_1/2_ is calculated using the formula: 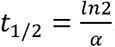, then is expressed as a fraction of total tree height.

### Bayes factor comparison of single vs. multi-optimum OU models

To evaluate whether the evolution of each trait is better described by adaptation toward multiple optima or a single global optimum, we fitted and compared alternative OU models using the Bayesian framework implemented in the R package “bayou”^41^ (version 2.3.2).

First, we fitted a multi-optimum OU model (OUM) that allows adaptive optima to shift across the phylogeny. Under this model, the strength of stabilizing evolution (*α*), the rate of stochastic trait evolution (*σ^2^*), the number of selective regimes (*k*), and the locations and values of adaptive optima (*θ*) were estimated jointly using a reversible-jump Markov chain Monte Carlo (rjMCMC) algorithm. We assigned half-Cauchy priors on both *α* and *σ^2^* (scale = 0.1), while the number of regime shifts was modelled using a conditional Poisson prior (*λ* = 1; maximum *k* = 50). The optima were assigned a normal prior centred on the empirical mean of the trait with a standard deviation equal to 1.5 times the trait’s observed standard deviation.

In parallel, we fitted a single-optimum OU model (OU1) for each trait to represent evolution toward a single global adaptive peak. In this model, *α* and *σ^2^* were estimated under the same half-Cauchy priors as in the OUM analyses, but the number of regime shifts was fixed to zero (*k* = 1), enforcing a single adaptive optimum across the entire phylogeny. For each trait, MCMC chains for both models (OU1 vs. OUM) were run for 100,000 iterations, with the first 30% of samples discarded as burn-in to ensure convergence, allowing for direct comparison between the models.

Model support was evaluated using Bayes factors based on marginal likelihoods estimated via stepping-stone sampling^91^. For each trait and model, marginal likelihoods were computed using 10,000 stepping-stone iterations. Log Bayes factors were calculated as the difference between the log marginal likelihoods of the OUM and OU1 models (*logBF* = *ln P*(*data* ∣ *OUM*) − *ln P*(*data* ∣ *OU1*)). Positive log Bayes factor values indicate support for the multi-optimum OUM model, whereas negative values indicate support for the single-optimum OU1 model. For the OUM models, posterior distributions of *k* were summarized using means and 95% highest posterior density (HPD) intervals.

### Definition of adaptive regimes

For growth rate, the mean estimated number of selective regimes across MCMC chains was 2.04, based on previous rjMCMC analyses, supporting the presence of approximately two growth-rate regimes across the phylogeny (Table S2). Therefore, we identified the node with the highest posterior probability of a regime shift and used it to define two distinct selective regimes: a slow-growing regime and a fast-growing regime (Figure 3b; Figure S2; Table S3). For other traits, we similarly defined two adaptive regimes per trait to simplify downstream coevolutionary analyses: the root regime and the regime corresponding to the node with the highest posterior probability of a regime shift (Figure S3; Table S3).

### Process-based evolutionary modelling

#### (1) Overview of the model

The multivariate Ornstein–Uhlenbeck (mvOU) process extends the univariate OU process^46,92^, which describes adaptive evolution of a trait toward an optimal value. The univariate OU process is parameterized by two key components: the selection strength (*α*) governing the rate of adaptation, and the optimum trait value (*θ*). In its multivariate form, this framework models the joint evolution of a *k*-dimensional suite of traits, 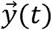, over a period of time through the following stochastic differential equation:

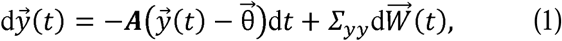

where 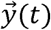 is the vector of trait values at time *t*; *A* is the matrix counterpart of the selection strength parameter *α*, represents rates of adaptation of trait values towards optimum values; 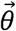 is the vector representing the optimum values for these traits; 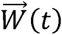 is a *k*-dimensional standard Brownian motion. The diffusion matrix *Σ_yy_* scales the resulting stochastic perturbations, which represent non-directed changes in trait values attributable to unconsidered selective factors, environmental fluctuations, or evolutionary constraints^42,45,93,94^.

To fully understand the coevolutionary dynamics of these traits, it is essential to examine both the *A* matrix directly, and to study its eigenvalues and eigenvectors^42,94^. The diagonal entries of *A*(*A_ii_*) represent the rate at which trait *i* returns to its own optimum *θ_i_*. The off-diagonal entries *A*(*A_ij_*) quantify coevolutionary coupling between traits, indicating how a deviation in trait *j* pulls trait *i* toward to (*A_ij_* > 0) or away from (*A_ij_* > 0) its optimum *θ_i_* (Figure S9).

Since *A* is the coefficient matrix in the OU stochastic differential equation (Eq. 1), each entry of this matrix captures the instantaneous rate of adaption/coevolution. In contrast, the eigenvalues (*λ*) of *A* describe the relative long-term rates at which linear combinations of traits evolve toward their optima along the independent directions in trait space defined by the corresponding eigenvectors 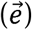 of *A*^42,94^ (Figure S9). Each eigenvector represents an independent evolutionary mode (direction) in trait space, with larger eigenvalues indicating faster evolution along those trajectories (Figure S9). To facilitate interpretation, we also reported the evolutionary half-life 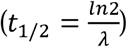 for each eigenvector. This metric represents the time required for an eigenvector to evolve halfway from its ancestral state to the optimum and serves as a measure of phylogenetic signal^95^. For clarity, we also report this half-life as a relative percentage of total tree height.

#### (2) Fitting single-regime bivariate OU models

To understand the tempo and mode of coevolution between growth rate and each associated trait, we fitted bivariate OU models in which both traits evolve toward constant, trait-specific optima (OU1). For comparison, we also fitted corresponding bivariate Brownian motion models (BM1) that assume neutral, unconstrained evolution. All model fitting was conducted using the R package *“mvSLOUCH”* (version 2.7.7), and model support was evaluated using AICc^96^, which has been shown to outperform other criteria (e.g., AIC, BIC) for differentiating among various OU model setups^42^. The relative fit of OU1 versus BM1 allowed us to determine whether coevolution between growth rate and each trait is better explained by stabilising evolution toward adaptive optima or by neutral drift. The phylogenetic tree was scaled to a total height of 1 to improve numerical stability and ensure more reliable parameter estimation when fitting all mvOU models, here and throughout.

We further explored the structure of coevolution by evaluating alternative parameterizations of the adaptation matrix (*A*) and the evolutionary diffusion matrix (*Σ*) to identify the best-supported representation of coevolutionary dynamics between growth rate and each trait over instantaneous timescales. Specifically, we compared models with different forms of the adaptation matrix (*A*-type = “*Any*,” “*Diagonal*,” “*UpperTri*,” “*LowerTri*”), which capture distinct hypotheses about the structure and directionality of adaptive interactions (Figure 1c). The “*Diagonal*” model assumes independent adaptation of each trait toward its own optimum, implying no coevolution. The “*UpperTri*” and “*LowerTri*” triangular models allow unidirectional evolutionary coupling, representing scenarios in which one trait responds adaptively to deviations in the other. The “*Any*” model provides a fully parameterized (asymmetric) adaptation matrix, allowing bidirectional co-adaptation between traits (Figure 1c). We also compared models with alternative parameterizations of the diffusion matrix (*Σ* = “*Diagonal*,” “*UpperTri*”) to test whether stochastic evolutionary drift occurs independently (“*Diagonal*”) or is correlated (“*UpperTri*”) between traits. Conclusions regarding the structure and directionality of coevolution were drawn from the best-supported model (i.e., model with the lowest AICc) for each growth rate – trait pair (Table S5).

#### (3) Fitting multi-regime bivariate OU models

To test whether distinct selective regimes impose different selection pressures on the coevolution between growth rate and each associated trait, we fitted multi-regime bivariate OU models (OUM). These models allow paired traits to evolve toward distinct adaptive optima under different regimes, thereby capturing directional shifts in selection across environments or lineages^36^.

For each trait pair, we evaluated whether coevolutionary dynamics were best explained by regimes defined by (i) the growth-rate regime, (ii) the trait-specific regime, or (iii) the combined regime integrating both growth rate and the trait. This framework allowed us to distinguish whether the coevolutionary relationship between growth rate and each trait was primarily structured by shifts in growth-rate evolution, by shifts in the associated trait, or by joint shifts in both traits.

#### (4) Fitting mvOU models with measurement error

Because multivariate OU models can be sensitive to uncertainty in trait estimates, we further assessed the robustness of model inference by incorporating alternative measurement-error settings^43,44^. For each trait pair, models were fitted under four measurement-error scenarios: (i) no measurement error, (ii) a low level of common measurement error (= 0.01) for all species, (iii) a medium level of common measurement error (= 0.05), and (iv) a high level of common measurement error (= 0.10; Figure S5; Table S4). These settings were used to evaluate whether support for particular evolutionary modes remained stable under increasing levels of observational uncertainty. The best-supported model across measurement-error settings was then used to summarize the dominant coevolutionary pattern for each growth-rate – trait pair (Table S5).

#### (5) Model optimization

To ensure that model comparison was based on well-converged solutions, parameter estimation for each mvOU model was conducted using a two-step procedure. First, we fitted a mvOU model in which the *A* matrix was specified as *“DecomposablePositive”* to obtain initial parameter estimates. Second, these estimates were used as starting values for a constrained optimization in which the structure of the *A* matrix was explicitly restricted according to the evolutionary mode being tested. This two-step procedure was repeated for 100 iterations using different random seeds. For each model, the solution with the lowest AICc from the second optimization step across all replicates was retained as the final result^42,94^.

## Author contribution

Study design: C.H. and N.K.; Data collection: C.H.; Methodology: C.H. and N.K.; Analysis: C.H.; Visualization: C.H.; Funding acquisition: C.H. and N.K.; Supervision: F.H.; Writing—original draft: C.H.; Writing—review and editing: C.H., N.K., Y.Z. and F.H.

## Acknowledgements

We thank Stuart West, Zhijie Liao, Qin li, Miaoxiao Wang and Xiaoyuan Feng for helpful discussions. We thank the National Natural Science Foundation of China (32602311: C.H.) and the Japan Society for the Promotion of Science (KAKENHI 24KJ0989: N.K.) for funding.

## Code and Data Availability

The data and code supporting the findings of this study will be made publicly available upon publication through a GitHub repository (https://github.com/haochh/coevo_growth). Intermediate results and model outputs necessary to reproduce the main analyses will also be provided. The original data were obtained from publicly available resources, as detailed in the Methods section and associated references.

## Use of generative AI

Generative artificial intelligence was used to assist with language editing and refinement of the manuscript for clarity and readability. All AI-assisted revisions were reviewed and approved by the authors, who take full responsibility for the content of the manuscript.

## References

1. Stewart, P. S. Growth rates of bacteria *in vivo*. Trends in Microbiology 33, 1056–1063 (2025).

2. Valiei, A., Dickson, A. M., Aminian-Dehkordi, J. & Mofrad, M. R. K. Bacterial community dynamics as a result of growth-yield trade-off and multispecies metabolic interactions toward understanding the gut biofilm niche. BMC Microbiol 24, 441 (2024).

3. Foley, M. M. et al. Growth rate as a link between microbial diversity and soil biogeochemistry. Nat Ecol Evol 8, 2018–2026 (2024).

4. He, X. et al. Microbial growth rate is a stronger predictor of soil organic carbon than carbon use efficiency. Nat Ecol Evol 10, 372–381 (2026).

5. Walker, T. W. N. et al. Microbial temperature sensitivity and biomass change explain soil carbon loss with warming. Nature Clim Change 8, 885–889 (2018).

6. Simon, E. et al. Microbial growth and carbon use efficiency show seasonal responses in a multifactorial climate change experiment. Commun Biol 3, 584 (2020).

7. Metze, D. et al. Microbial growth under drought is confined to distinct taxa and modified by potential future climate conditions. Nat Commun 14, 5895 (2023).

8. Zhou, Z. et al. The biogeography of soil microbiome potential growth rates. Nat Commun 15, 9472 (2024).

9. López, J. L. et al. Growth rate is a dominant factor predicting the rhizosphere effect. ISME J 17, 1396–1405 (2023).

10. Levin, P. A. & Wang, J. D. Bacterial growth rate in populations and single cells. Nat Microbiol 1–9 (2026) doi:10.1038/s41564-026-02449-6.

11. Thiermann, R. et al. Decoupling of global metabolic flux and proteome partitioning in bacteria. Science 392, eaeb6410 (2026).

12. Gibson, B., Wilson, D. J., Feil, E. & Eyre-Walker, A. The distribution of bacterial doubling times in the wild. Proc Biol Sci 285, 20180789 (2018).

13. Weissman, J. L., Hou, S. & Fuhrman, J. A. Estimating maximal microbial growth rates from cultures, metagenomes, and single cells via codon usage patterns. Proceedings of the National Academy of Sciences 118, e2016810118 (2021).

14. Deulofeu-Capo, O. et al. Growth rates of marine prokaryotes are extremely diverse, even among closely related taxa. ISME Communications 4, ycae066 (2024).

15. Xu, L., Zakem, E. & Weissman, J. L. Improved maximum growth rate prediction from microbial genomes by integrating phylogenetic information. Nat Commun 16, 4256 (2025).

16. Sezonov, G., Joseleau-Petit, D. & D’Ari, R. Escherichia coli Physiology in Luria-Bertani Broth. Journal of Bacteriology 189, 8746–8749 (2007).

17. Testerman, T. L., McGee, D. J. & Mobley, H. L. T. Helicobacter pylori Growth and Urease Detection in the Chemically Defined Medium Ham’s F-12 Nutrient Mixture. Journal of Clinical Microbiology 39, 3842–3850 (2001).

18. Starnawski, P. et al. Microbial community assembly and evolution in subseafloor sediment. Proceedings of the National Academy of Sciences 114, 2940–2945 (2017).

19. Trembath-Reichert, E. et al. Methyl-compound use and slow growth characterize microbial life in 2-km-deep subseafloor coal and shale beds. Proceedings of the National Academy of Sciences 114, E9206–E9215 (2017).

20. Hessen, D. O., Jeyasingh, P. D., Neiman, M. & Weider, L. J. Genome streamlining and the elemental costs of growth. Trends in Ecology & Evolution 25, 75–80 (2010).

21. Lynch, M. Streamlining and Simplification of Microbial Genome Architecture. Annual Review of Microbiology 60, 327–349 (2006).

22. Giovannoni, S. J., Cameron Thrash, J. & Temperton, B. Implications of streamlining theory for microbial ecology. ISME J 8, 1553–1565 (2014).

23. Vieira-Silva, S. & Rocha, E. P. C. The Systemic Imprint of Growth and Its Uses in Ecological (Meta)Genomics. PLOS Genetics 6, e1000808 (2010).

24. Mira, A., Ochman, H. & Moran, N. A. Deletional bias and the evolution of bacterial genomes. Trends in Genetics 17, 589–596 (2001).

25. Touchon, M. & Rocha, E. P. C. Causes of Insertion Sequences Abundance in Prokaryotic Genomes. Molecular Biology and Evolution 24, 969–981 (2007).

26. Grafen, A. & Hamilton, W. D. The phylogenetic regression. Philosophical Transactions of the Royal Society of London. B, Biological Sciences 326, 119–157 (1989).

27. Hadfield, J. D. MCMC Methods for Multi-Response Generalized Linear Mixed Models: The MCMCglmm R Package. Journal of Statistical Software 33, 1–22 (2010).

28. Ringen, E. J., Claessens, S., Martin, J. S. & Jaeggi, A. V. Trait coevolution and causal inference using generalized dynamic phylogenetic models. https://besjournals.onlinelibrary.wiley.com/doi/10.1111/2041-210x.70303.

29. Hansen, T. F. STABILIZING SELECTION AND THE COMPARATIVE ANALYSIS OF ADAPTATION. Evol 51, 1341–1351 (1997).

30. Butler, M. A. & King, A. A. Phylogenetic Comparative Analysis: A Modeling Approach for Adaptive Evolution. The American Naturalist 164, 683–695 (2004).

31. Bartoszek, K., Pienaar, J., Mostad, P., Andersson, S. & Hansen, T. F. A phylogenetic comparative method for studying multivariate adaptation. Journal of Theoretical Biology 314, 204–215 (2012).

32. Pienaar, J. et al. Phylogenetic comparative methods for studying adaptation: the adaptation-inertia framework. *j. evol*. Biol. 39, 1–17 (2026).

33. Zhang, Y. et al. Multivariate Coevolution Shapes Life-History Strategies Across Amniotes. 2025.03.09.642183 Preprint at 10.1101/2025.03.09.642183 (2026).

34. Price, P. D. et al. Detecting signatures of selection on gene expression. Nat Ecol Evol 6, 1035–1045 (2022).

35. Cope, A. L., Schraiber, J. G. & Pennell, M. Macroevolutionary divergence of gene expression driven by selection on protein abundance. Science 387, 1063–1068 (2025).

36. Sol, D. et al. Adaptations to marine environments and the evolution of slow-paced life histories in endotherms. Nat Commun 16, 4265 (2025).

37. Neupane, S., Zanne, A. E., Lens, F. & Uyeda, J. C. Modeling trait heterogeneity and inferring causal links in the macroevolution of growth habit in eudicot angiosperms. New Phytologist n/a,.

38. Madin, J. S. et al. A synthesis of bacterial and archaeal phenotypic trait data. Sci Data 7, 170 (2020).

39. Blomberg, S. P., Garland JR., T. & Ives, A. R. Testing for Phylogenetic Signal in Comparative Data: Behavioral Traits Are More Labile. Evolution 57, 717–745 (2003).

40. Freckleton, R. P., Harvey, P. H. & Pagel, M. Phylogenetic Analysis and Comparative Data: A Test and Review of Evidence. The American Naturalist 160, 712–726 (2002).

41. Uyeda, J. C. & Harmon, L. J. A Novel Bayesian Method for Inferring and Interpreting the Dynamics of Adaptive Landscapes from Phylogenetic Comparative Data. Syst Biol 63, 902–918 (2014).

42. Bartoszek, K. et al. Fast mvSLOUCH: Multivariate Ornstein–Uhlenbeck-based models of trait evolution on large phylogenies. Methods in Ecology and Evolution 15, 1507–1515 (2024).

43. Silvestro, D., Kostikova, A., Litsios, G., Pearman, P. B. & Salamin, N. Measurement errors should always be incorporated in phylogenetic comparative analysis. Methods in Ecology and Evolution 6, 340–346 (2015).

44. Cooper, N., Thomas, G. H., Venditti, C., Meade, A. & Freckleton, R. P. A cautionary note on the use of Ornstein Uhlenbeck models in macroevolutionary studies. Biol J Linn Soc 118, 64–77 (2016).

45. Bartoszek, K., Pienaar, J., Mostad, P., Andersson, S. & Hansen, T. F. A phylogenetic comparative method for studying multivariate adaptation. Journal of Theoretical Biology 314, 204–215 (2012).

46. Hansen, T. F. Stabilizing selection and the comparative analysis of adaptation. Evolution 51, 1341–1351 (1997).

47. Cornwallis, C. K. & Griffin, A. S. A Guided Tour of Phylogenetic Comparative Methods for Studying Trait Evolution. *Annual Review of Ecology*, Evolution, and Systematics 55, 181–204 (2024).

48. Zhu, M. & Dai, X. Shaping of microbial phenotypes by trade-offs. Nat Commun 15, 4238 (2024).

49. Björkeroth, J. et al. Proteome reallocation from amino acid biosynthesis to ribosomes enables yeast to grow faster in rich media. Proceedings of the National Academy of Sciences 117, 21804–21812 (2020).

50. Belliveau, N. M. et al. Fundamental limits on the rate of bacterial growth and their influence on proteomic composition. cels 12, 924–944.e2 (2021).

51. Scott, M. & Hwa, T. Shaping bacterial gene expression by physiological and proteome allocation constraints. Nat Rev Microbiol 21, 327–342 (2023).

52. Zhu, M., Mori, M., Hwa, T. & Dai, X. Distantly related bacteria share a rigid proteome allocation strategy with flexible enzyme kinetics. Proceedings of the National Academy of Sciences 122, e2427091122 (2025).

53. Ho, A., Di Lonardo, D. P. & Bodelier, P. L. E. Revisiting life strategy concepts in environmental microbial ecology. FEMS Microbiol Ecol 93, fix006 (2017).

54. Stone, B. W. G. et al. Life history strategies among soil bacteria—dichotomy for few, continuum for many. The ISME Journal 17, 611–619 (2023).

55. Hernandez, D. J., Kiesewetter, K. N., Almeida, B. K., Revillini, D. & Afkhami, M. E. Multidimensional specialization and generalization are pervasive in soil prokaryotes. Nat Ecol Evol 7, 1408–1418 (2023).

56. Carscadden, K. A. et al. Niche Breadth: Causes and Consequences for Ecology, Evolution, and Conservation. The Quarterly Review of Biology 95, 179–214 (2020).

57. Sexton, J. P., Montiel, J., Shay, J. E., Stephens, M. R. & Slatyer, R. A. Evolution of Ecological Niche Breadth. Annual Review of Ecology, Evolution, and Systematics 48, 183–206 (2017).

58. Into the microbial niche. Trends in Ecology & Evolution 38, 936–945 (2023).

59. Hao, C. et al. Cooperation and the evolution of bacterial niche breadth. Proceedings of the National Academy of Sciences 123, e2520645123 (2026).

60. Polz, M. F., Alm, E. J. & Hanage, W. P. Horizontal gene transfer and the evolution of bacterial and archaeal population structure. Trends in Genetics 29, 170–175 (2013).

61. Bobay, L.-M. & Ochman, H. The Evolution of Bacterial Genome Architecture. Front. Genet. 8, (2017).

62. Taylor, J. S. & Raes, J. Duplication and Divergence: The Evolution of New Genes and Old Ideas. Annual Review of Genetics 38, 615–643 (2004).

63. Treangen, T. J. & Rocha, E. P. C. Horizontal Transfer, Not Duplication, Drives the Expansion of Protein Families in Prokaryotes. PLOS Genetics 7, e1001284 (2011).

64. Gao, Y. & Wu, M. Microbial genomic trait evolution is dominated by frequent and rare pulsed evolution. Science Advances 8, eabn1916 (2022).

65. Roller, B. R. K., Stoddard, S. F. & Schmidt, T. M. Exploiting rRNA operon copy number to investigate bacterial reproductive strategies. Nat Microbiol 1, 16160 (2016).

66. Uyeda, J. C., Zenil-Ferguson, R. & Pennell, M. W. Rethinking phylogenetic comparative methods. Syst Biol 67, 1091–1109 (2018).

67. Ringen, E. J., Claessens, S., Martin, J. S. & Jaeggi, A. V. Trait coevolution and causal inference using generalized dynamic phylogenetic models. Methods in Ecology and Evolution **n/a**,.

68. Del Giudice, M. Rethinking the fast-slow continuum of individual differences. Evolution and Human Behavior 41, 536–549 (2020).

69. Van de Walle, J. et al. Individual life histories: neither slow nor fast, just diverse. Proceedings of the Royal Society B: Biological Sciences 290, 20230511 (2023).

70. Lu, M. et al. A theoretical framework for scaling ecological niches from individuals to species. Proceedings of the National Academy of Sciences 122, e2425582122 (2025).

71. Agrawal, A. A. A scale-dependent framework for trade-offs, syndromes, and specialization in organismal biology. Ecology 101, e02924 (2020).

72. Grabowski, M. et al. A Cautionary Note on “A Cautionary Note on the Use of Ornstein Uhlenbeck Models in Macroevolutionary Studies”. Syst Biol 72, 955–963 (2023).

73. Parks, D. H. et al. GTDB: an ongoing census of bacterial and archaeal diversity through a phylogenetically consistent, rank normalized and complete genome-based taxonomy. Nucleic Acids Research 50, D785–D794 (2022).

74. Bowers, R. M. et al. Minimum information about a single amplified genome (MISAG) and a metagenome-assembled genome (MIMAG) of bacteria and archaea. Nat Biotechnol 35, 725–731 (2017).

75. Benson, D. A. et al. GenBank. Nucleic Acids Research 41, D36–D42 (2013).

76. O’Leary, N. A. et al. Reference sequence (RefSeq) database at NCBI: current status, taxonomic expansion, and functional annotation. Nucleic Acids Research 44, D733–D745 (2016).

77. Roller, B. R. K., Stoddard, S. F. & Schmidt, T. M. Exploiting rRNA operon copy number to investigate bacterial reproductive strategies. Nat Microbiol 1, 1–7 (2016).

78. Chen, Y.-J. et al. Metabolic flexibility allows bacterial habitat generalists to become dominant in a frequently disturbed ecosystem. ISME J 15, 2986–3004 (2021).

79. D’Souza, G. et al. Ecology and evolution of metabolic cross-feeding interactions in bacteria. Natural Product Reports 35, 455–488 (2018).

80. Ramoneda, J., Jensen, T. B. N., Price, M. N., Casamayor, E. O. & Fierer, N. Taxonomic and environmental distribution of bacterial amino acid auxotrophies. Nat Commun 14, 7608 (2023).

81. Kanehisa, M., Sato, Y., Kawashima, M., Furumichi, M. & Tanabe, M. KEGG as a reference resource for gene and protein annotation. Nucleic Acids Research 44, D457–D462 (2016).

82. Aramaki, T. et al. KofamKOALA: KEGG Ortholog assignment based on profile HMM and adaptive score threshold. Bioinformatics 36, 2251–2252 (2020).

83. Richardson, L. et al. MGnify: the microbiome sequence data analysis resource in 2023. Nucleic Acids Research 51, D753–D759 (2023).

84. Barnum, T. P. et al. Predicting microbial growth conditions from amino acid composition. 2024.03.22.586313 Preprint at 10.1101/2024.03.22.586313 (2024).

85. Lu, M., Winner, K. & Jetz, W. A unifying framework for quantifying and comparing n-dimensional hypervolumes. Methods in Ecology and Evolution 12, 1953–1968 (2021).

86. Sriswasdi, S., Yang, C. & Iwasaki, W. Generalist species drive microbial dispersion and evolution. Nat Commun 8, 1162 (2017).

87. Mise, K. & Iwasaki, W. Environmental Atlas of Prokaryotes Enables Powerful and Intuitive Habitat-Based Analysis of Community Structures. iScience 23, 101624 (2020).

88. Paradis, E. & Schliep, K. ape 5.0: an environment for modern phylogenetics and evolutionary analyses in R. Bioinformatics 35, 526–528 (2019).

89. Revell, L. J. phytools: an R package for phylogenetic comparative biology (and other things). Methods in Ecology and Evolution 3, 217–223 (2012).

90. Pennell, M. W. et al. geiger v2.0: an expanded suite of methods for fitting macroevolutionary models to phylogenetic trees. Bioinformatics 30, 2216–2218 (2014).

91. Xie, W., Lewis, P. O., Fan, Y., Kuo, L. & Chen, M.-H. Improving Marginal Likelihood Estimation for Bayesian Phylogenetic Model Selection. Syst Biol 60, 150–160 (2011).

92. Butler, M. A. & King, A. A. Phylogenetic Comparative Analysis: A Modeling Approach for Adaptive Evolution. (2004).

93. Clavel, J., Escarguel, G. & Merceron, G. mv morph : an r package for fitting multivariate evolutionary models to morphometric data. Methods Ecol Evol 6, 1311–1319 (2015).

94. Bartoszek, K. et al. Model Selection Performance in Phylogenetic Comparative Methods Under Multivariate Ornstein–Uhlenbeck Models of Trait Evolution. Systematic Biology 72, 275–293 (2023).

95. Hansen, T. F. & Bartoszek, K. Interpreting the Evolutionary Regression: The Interplay Between Observational and Biological Errors in Phylogenetic Comparative Studies. Systematic Biology 61, 413–425 (2012).

96. Hurvich, C. M. & Tsai, C.-L. Regression and Time Series Model Selection in Small Samples. Biometrika 76, 297–307 (1989).

