## Supplementary materials for "Growth rate structures the dynamics of bacterial trait evolution"

### Supplementary figures (S1 to S10) to support findings in the main text

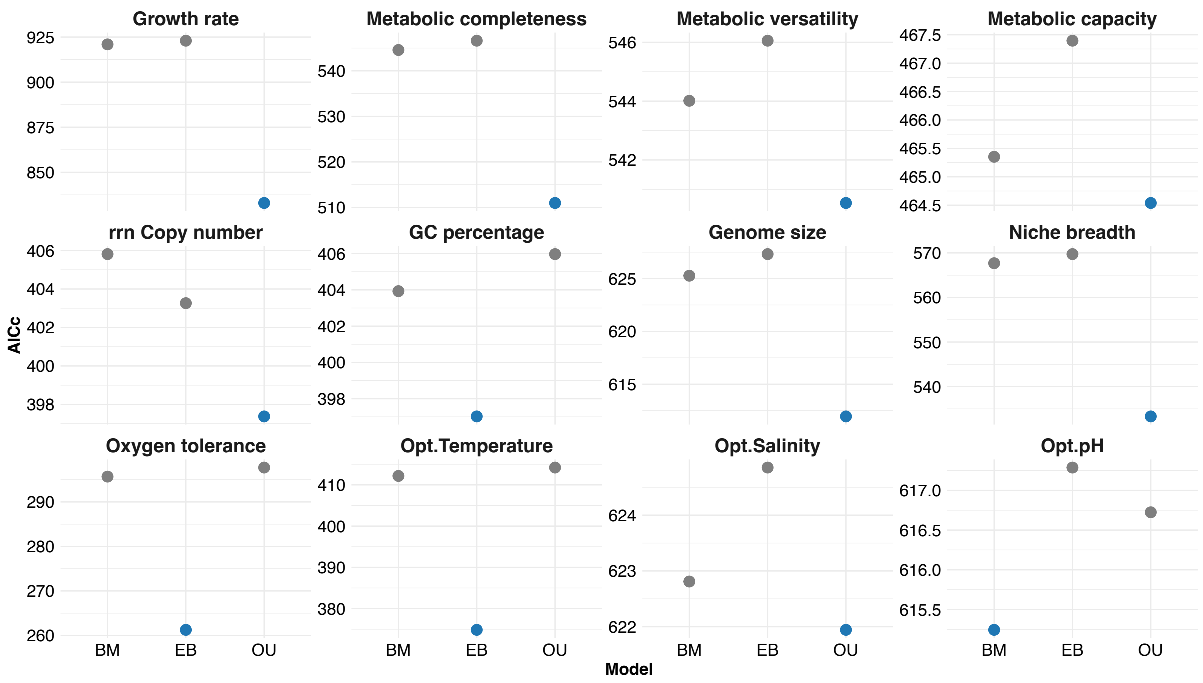

**Figure S1.** Comparison of goodness-of-fit for the evolutionary mode of each individual trait based on AICc among three commonly used trait evolution models: Brownian motion (BM), Ornstein–Uhlenbeck (OU), and early burst (EB). AICc values are plotted for each model across the traits, with lower AICc values indicating a better model fit. The blue dot represents the model that best explains each trait. The growth rate is best explained by the OU model.

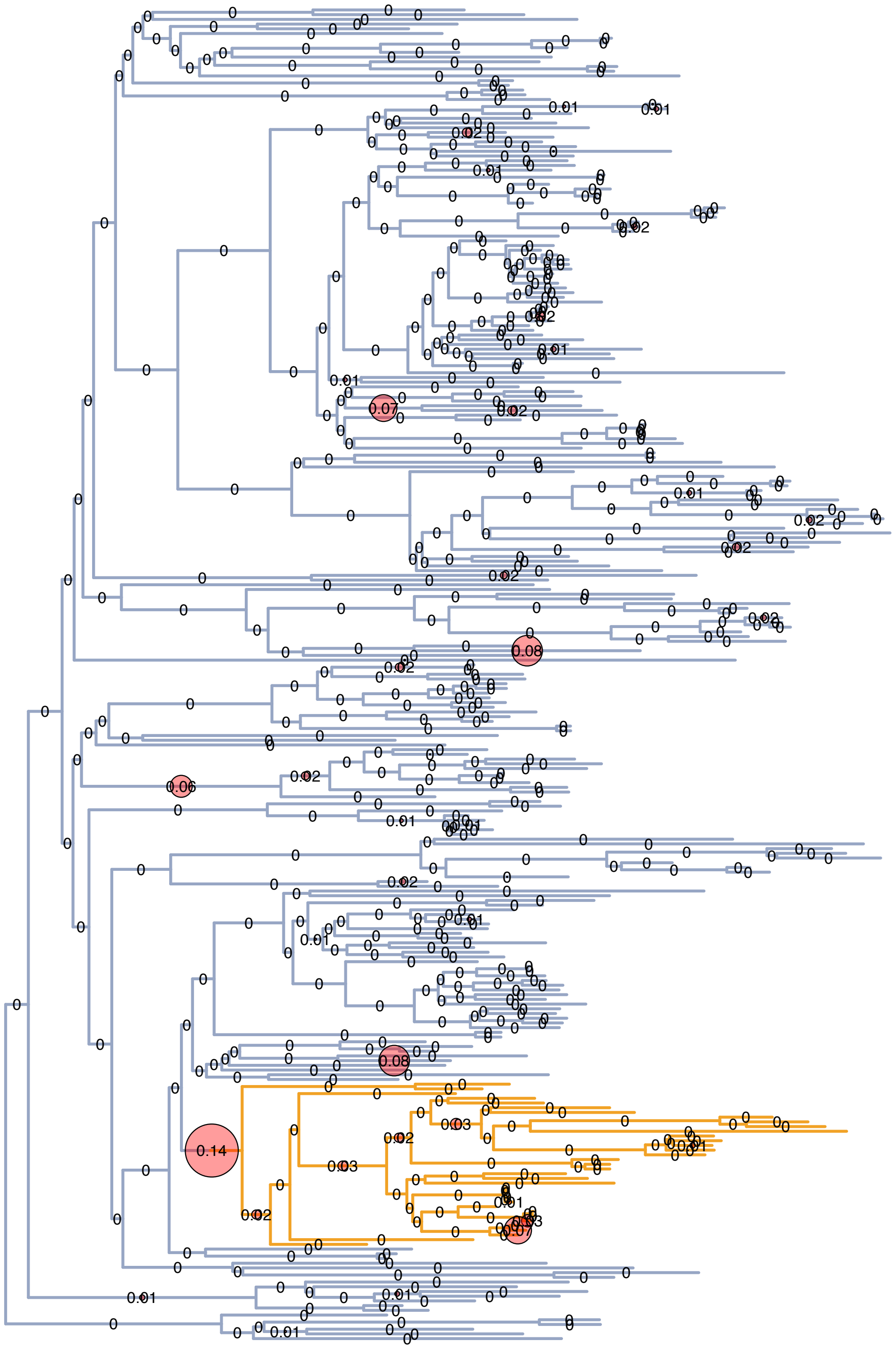

**Figure S2.** Phylogenetic distribution of selective regimes for growth rate inferred from a *bayou* reversible-jump MCMC analysis under a multi-regime OU model. Circles of different sizes indicate the posterior probability of a regime shift at the corresponding node. The node with the highest posterior probability of a regime shift was used to define two distinct selective regimes for growth rate: a slow-growing regime (the root regime, shown in blue) and a fast-growing regime (shown in orange).

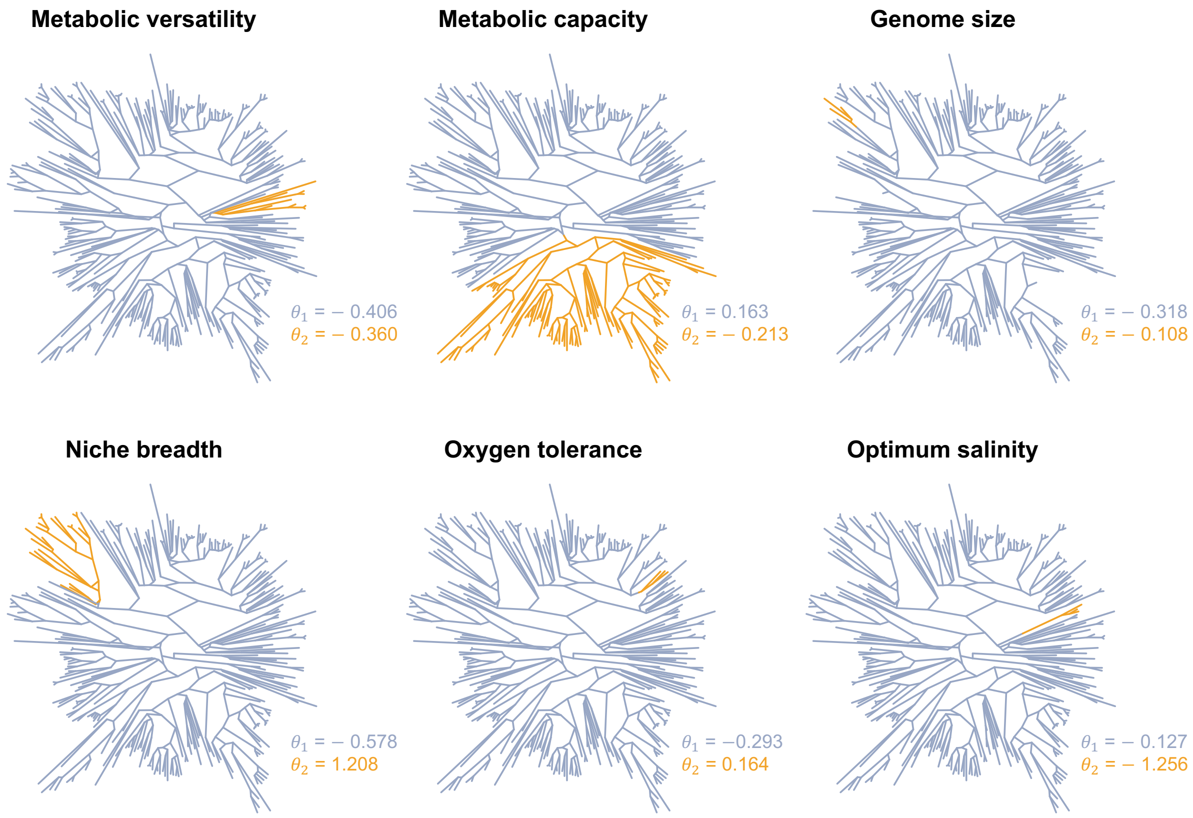

**Figure S3**. Phylogenetic distribution of selective regimes for the six traits for which Bayes factor comparisons favoured the OUM model over the OU1 model. For each trait, the root regime is shown in blue and the shifted regime is shown in orange. The corresponding adaptive optima ($\theta$) for each regime are mapped onto the phylogeny.

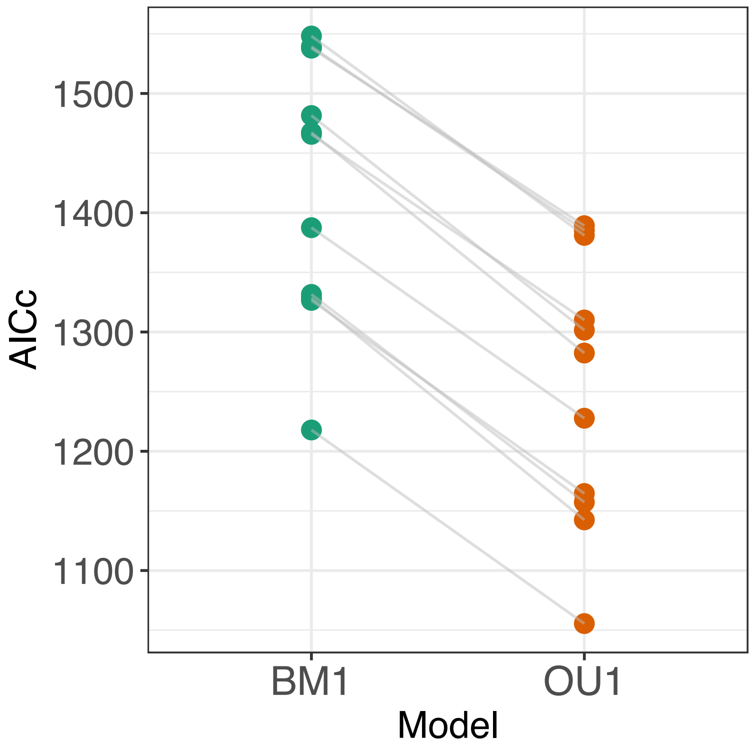

**Figure S4.** Comparison of model fit between Brownian motion and Ornstein–Uhlenbeck models for trait coevolution. Goodness-of-fit based on AICc was presented for the coevolution of growth rate and each of the 11 traits under a Brownian motion model (BM1) and a single-regime Ornstein–Uhlenbeck model (OU1). Lower AICc values indicate better model support. Across all trait pairs, OU1 models consistently show improved fit relative to BM1, supporting a model of coevolution driven by stabilizing evolution toward trait-specific optima rather than neutral drift.

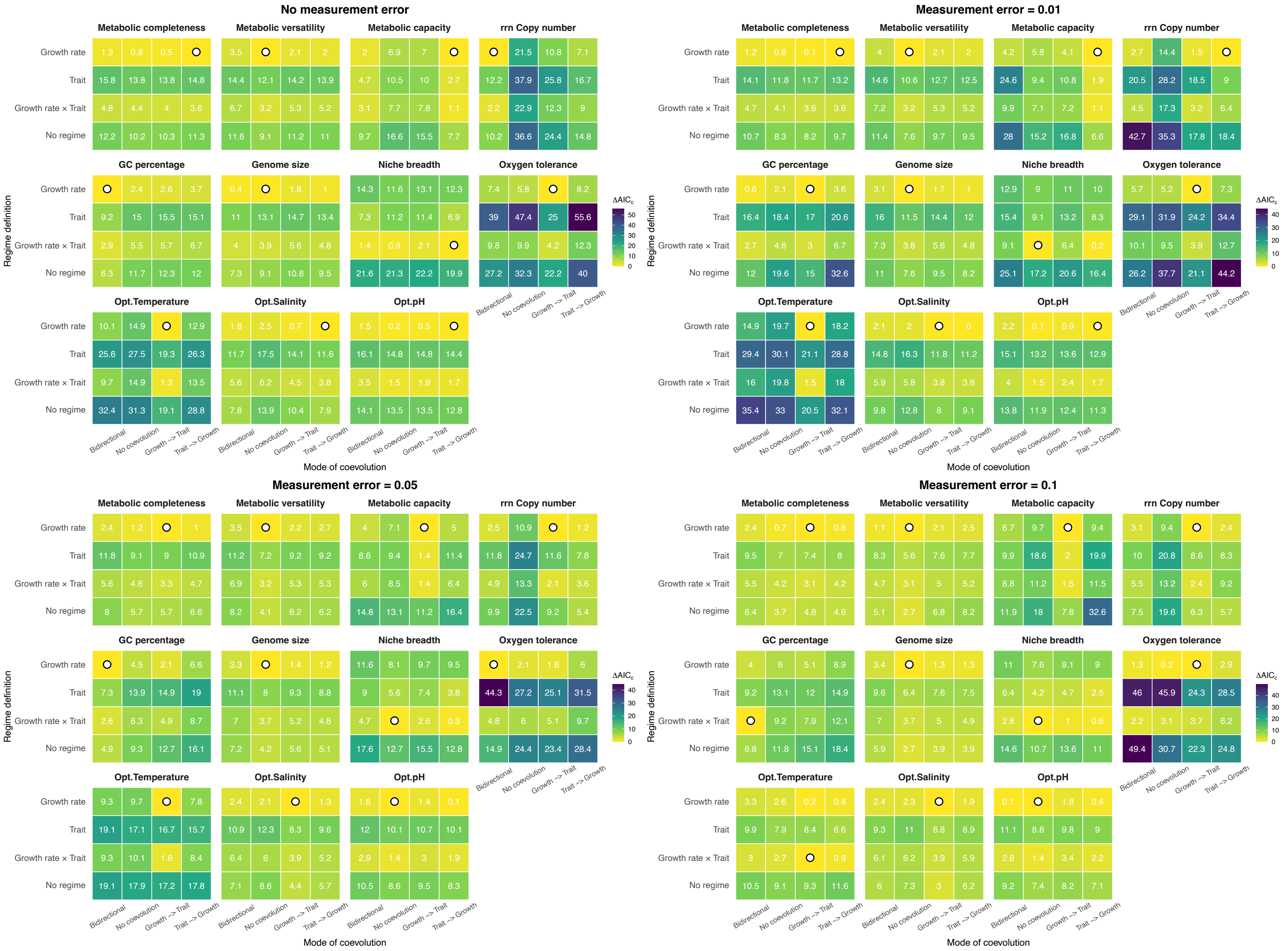

**Figure S5.** Coevolutionary modes between growth rate and associated traits across alternative measurement-error settings. Bivariate OU models were fitted for growth rate and each of the 11 associated traits under 4 measurement-error settings: (i) no measurement error, (ii) a low level of common measurement error (= 0.01) for all species, (iii) a medium level of common measurement error (= 0.05), and (iv) a high level of common measurement error (= 0.10). Each panel compares the goodness of fit of alternative evolutionary models for one growth-rate – trait pair. The x-axis indicates the presence or absence of coevolution and, where applicable, the inferred directionality of coevolutionary coupling. The y-axis indicates the selective-regime definition under which coevolution was evaluated. Heatmap colours represent ΔAICc values, with lower values closer to zero indicating better model fit. A circle within a grid cell indicates the best-supported model for that trait pair (ΔAICc = 0). Results are shown for models in which the matrix was constrained to a "*Diagonal*" structure, thereby assuming independent stochastic perturbations between the two traits.

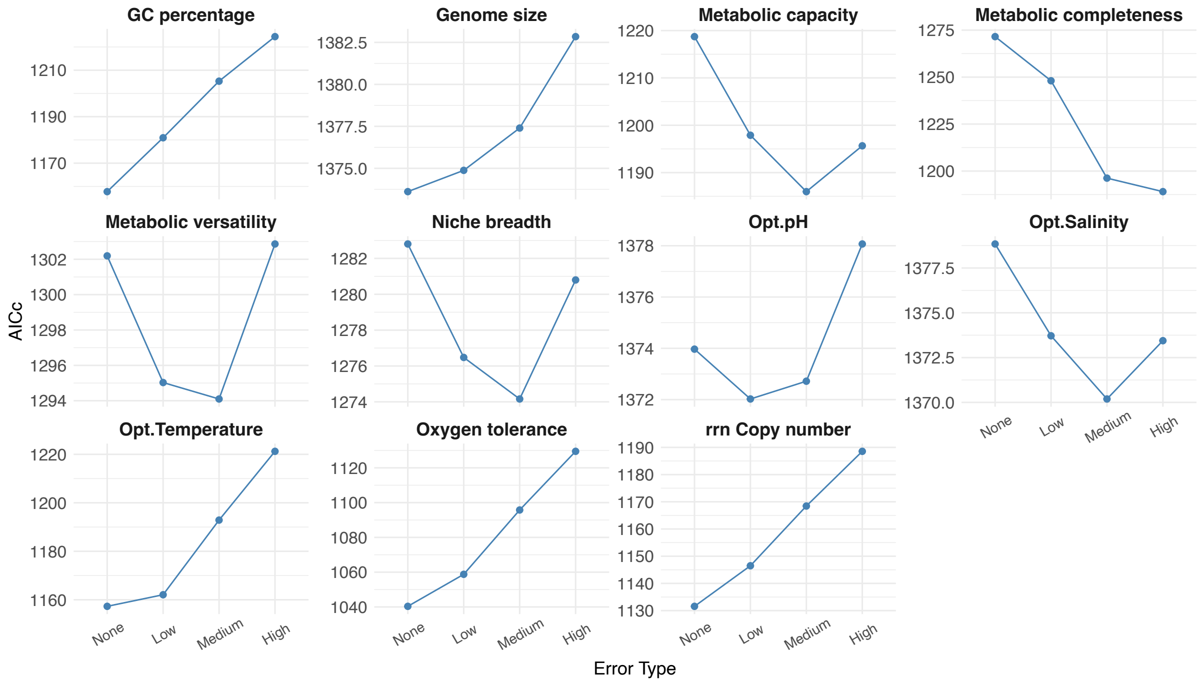

**Figure S6.** Comparison of goodness-of-fit based on AICc for coevolution between growth rate and associated traits across alternative measurement-error settings. Bivariate OU models were fitted for growth rate and each of the 11 traits under four measurement-error scenarios: (i) no measurement error, (ii) low common measurement error (= 0.01), (iii) medium common measurement error (= 0.05), and (iv) high common measurement error (= 0.10).

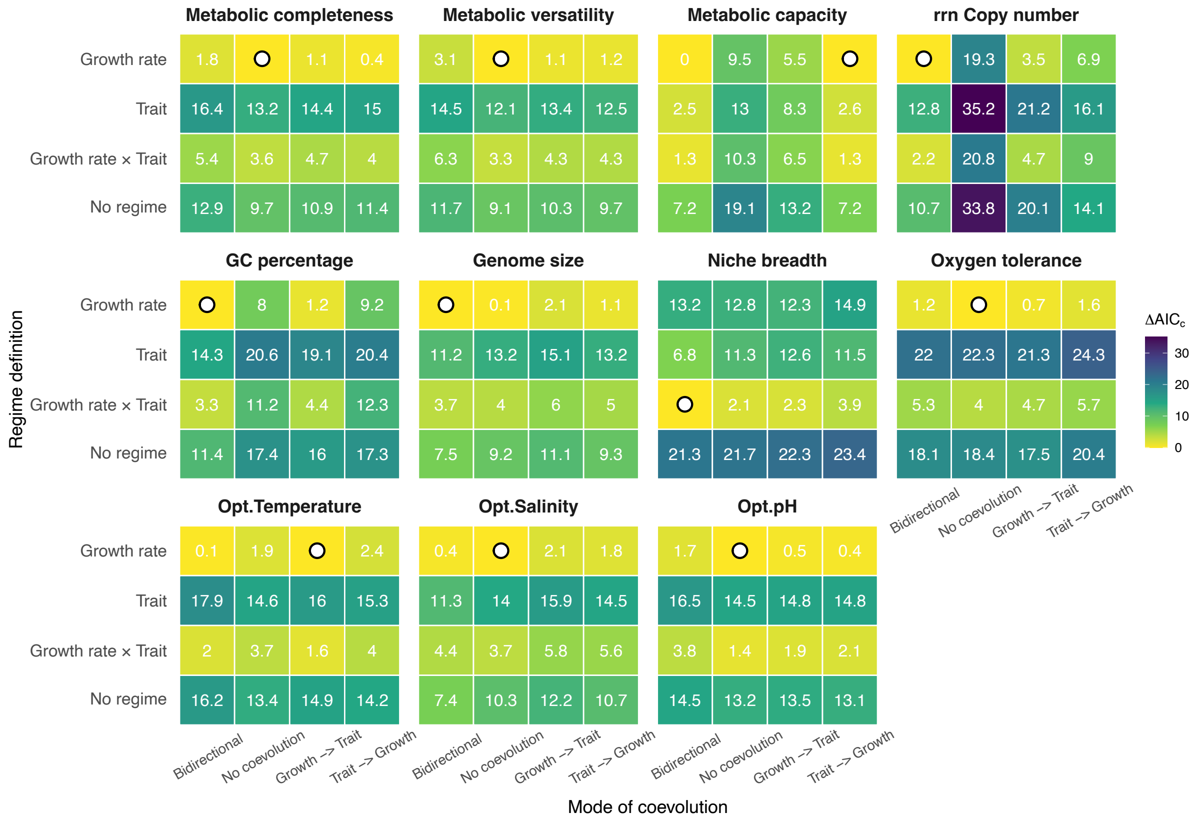

**Figure S7.** Mode of coevolution between growth rate and each trait when assuming a "*UpperTri*" structure for the $\Sigma_{yy}$ matrix, which treats the stochastic perturbations between the two traits as correlated. Coevolution was modelled without considering the measurement errors. Each panel represents the comparison of goodness-of-fit for bivariate OU models between growth rate and one of the 11 traits under different evolutionary modes. The x-axis shows the presence/absence and directionality of coevolution, while the y-axis represents the regime definitions under which coevolution is evaluated. The heatmap colours represent the ΔAICc values, with lower values (closer to zero) indicating better model fits. A circle inside a grid indicates the model with the best fit (ΔAICc = 0) for the coevolution between growth rate and that trait pair.

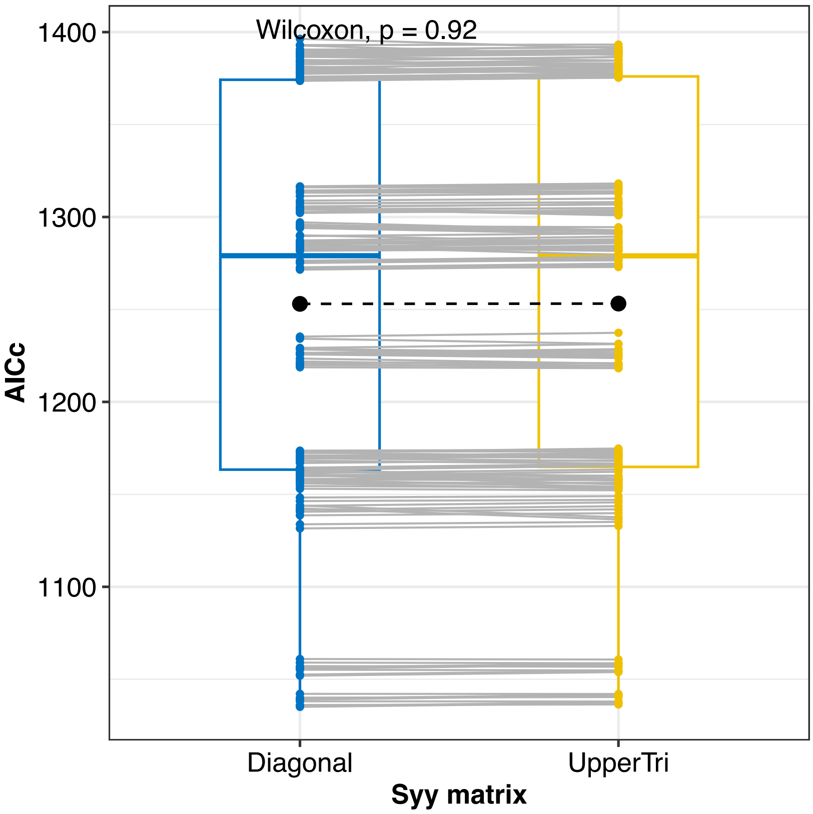

**Figure S8.** Comparison of model fit between $\Sigma_{yy}$ matrix assumptions for bivariate OU models of growth rate and each trait under two different assumptions for the $\Sigma_{yy}$ (stochastic perturbation) matrix: "*Diagonal*" (assuming independent perturbations between traits) and "*UpperTri*" (assuming correlated perturbations). Each line connects the AICc values for a single trait pair with the same evolutionary mode specification, which includes the presence/absence and directionality of coevolution as well as the regime definitions. The black dots indicate the mean AICc for each $\Sigma_{yy}$ type. No significant difference in model fit was observed between the two $\Sigma_{yy}$ assumptions (Wilcoxon test, p = 0.92).

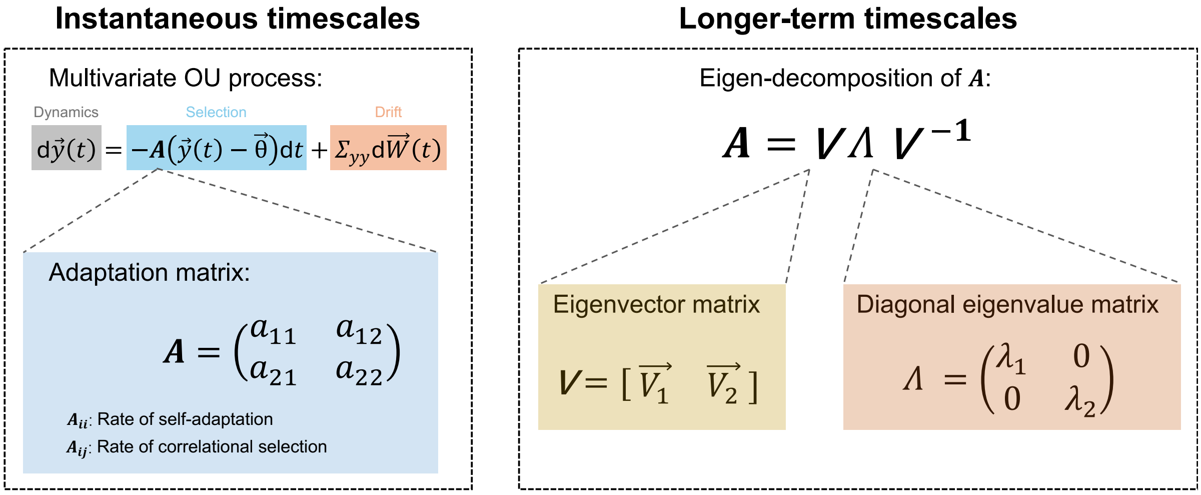

**Figure S9.** Rates of coevolution at instantaneous and longer-term timescales under a multivariate Ornstein–Uhlenbeck (mvOU) framework. Left panel: The multivariate OU process models the coevolution between two traits using a stochastic differential equation. The rate parameters in the adaptation matrix (***A*** matrix) represent the speed of adaptation of trait values over instantaneous timescales. Right panel: Eigen-decomposition of the ***A*** matrix allows examination of how linear combinations of traits evolve along independent adaptive axes over longer evolutionary timescales. Each eigenvector defines an axis of co-adaptation in the joint trait space, and the corresponding eigenvalue quantifies the strength of adaptive pull along that axis. Thus, the eigen-decomposition of the ***A*** matrix provides insight into longer-term coevolutionary dynamics between traits.

| | Metabolic completeness | | Metabolic versatility | | Metabolic capacity | | rrn Copy number | | GC percentage | | Genome size | |
|---|---|---|---|---|---|---|---|---|---|---|---|---|
| | $\vec{e1}$ | $\vec{e2}$ | $\vec{e1}$ | $\vec{e2}$ | $\vec{e1}$ | $\vec{e2}$ | $\vec{e1}$ | $\vec{e2}$ | $\vec{e1}$ | $\vec{e2}$ | $\vec{e1}$ | $\vec{e2}$ |
| Growth rate | 0.984 | 0.000 | -1.000 | 0.000 | 0.979 | 0.000 | 0.982 | 0.460 | 0.984 | -0.238 | -1.000 | 0.000 |
| Factor | -0.180 | 1.000 | 0.000 | -1.000 | 0.203 | 1.000 | 0.187 | 0.888 | -0.179 | 0.971 | 0.000 | -1.000 |
| Eigenvalue ($\lambda$) | 5.965 | 0.157 | 7.750 | 0.342 | 7.868 | 0.012 | 26.937 | 2.811 | 33.573 | 1.147 | 15.116 | 1.988 |
| Half-life ($t_{1/2} = \frac{ln2}{\lambda}$) | 0.116 | 4.409 | 0.089 | 2.027 | 0.088 | 57.233 | 0.026 | 0.247 | 0.021 | 0.604 | 0.046 | 0.349 |
| Tree Height (%) | 11.620 | 440.850 | 8.944 | 202.658 | 8.810 | 5723.257 | 2.573 | 24.655 | 2.065 | 60.439 | 4.585 | 34.862 |

| | Niche breadth | | Oxygen tolerance | | Opt. Temperature | | Opt. Salinity | | Opt. pH | |
|---|---|---|---|---|---|---|---|---|---|---|
| | $\vec{e1}$ | $\vec{e2}$ | $\vec{e1}$ | $\vec{e2}$ | $\vec{e1}$ | $\vec{e2}$ | $\vec{e1}$ | $\vec{e2}$ | $\vec{e1}$ | $\vec{e2}$ |
| Growth rate | -1.000 | 0.000 | 0.992 | 0.000 | 0.977 | 0.000 | 0.979 | 0.000 | 1.000 | -0.134 |
| Factor | 0.000 | -1.000 | -0.126 | 1.000 | -0.211 | 1.000 | -0.205 | 1.000 | 0.000 | 0.991 |
| Eigenvalue ($\lambda$) | 7.338 | 2.629 | 17.540 | 0.026 | 24.239 | 1.118 | 7.643 | 0.460 | 12.491 | 0.286 |
| Half-life ($t_{1/2} = \frac{ln2}{\lambda}$) | 0.094 | 0.264 | 0.040 | 26.636 | 0.029 | 0.620 | 0.091 | 1.507 | 0.055 | 2.420 |
| Tree Height (%) | 9.445 | 26.364 | 3.952 | 2663.619 | 2.860 | 61.985 | 9.069 | 150.697 | 5.549 | 242.003 |

**Figure S10.** Summary of the key parameters describing the coevolutionary dynamics between growth rate and each associated trait over longer timescales, derived from the eigen-decomposition of the ***A*** matrices of the best-supported models. For each trait pair, the table lists:

(1) eigenvectors ($\vec{e_1}, \vec{e_2}$): each eigenvector represent a main axis of evolutionary change over longer evolutionary timescales. For each eigenvector, loadings of growth rate and the focal trait along the two primary axes were presented, indicating the combination of traits that evolve together alongside the direction defined by that eigenvector. Positive and negative values denote whether traits change in the same or opposite directions along that axis.

(2) eigenvalues ($\lambda_1, \lambda_2$): associated rates of evolution along each eigenvector, reflecting the speed of evolutionary change.

(3) evolutionary half-lives ($t_{1/2}$): calculated as $\frac{ln2}{\lambda}$, representing the timescale required for a trait combination to evolve halfway toward its optimum, expressed as a fraction of total tree height.

#### Supplementary analysis 1: Evolutionary mode of bioinformatically estimated growth rate

Our primary analyses were based on experimentally measured maximum growth rates to prioritize data accuracy. However, a limitation of this approach is that experimentally measured growth-rate data are available for a relatively small number of species in our dataset (283 species), compared with the known diversity of bacteria[1]. We therefore performed a supplementary analysis using a larger dataset of bioinformatically estimated growth rates. Although these estimates are less direct than experimental measurements and may contain additional uncertainty, they provide broader taxonomic coverage and allow us to evaluate whether similar evolutionary patterns are detectable across a larger set of species. Specifically, we used this expanded dataset to characterize the evolutionary mode of growth rate and 11 associated traits.

##### Species selection and genome collection

We obtained bacterial species and their representative genomes from the Genome Taxonomy Database (GTDB[2]; release 220) on 1st July 2024. Species were selected if their representative genomes allowed identification of full-length 16S rRNA gene sequences. For these species, we adopted a stricter standard than the Minimum Information about a Metagenome-Assembled Genome (MIMAG) criterion[3], retaining only high-quality genomes with completeness ≥ 95% and contamination ≤ 5%. Nucleotide and amino acid sequences for all representative genomes were downloaded using *ncbi-genome-download* scripts (version 0.3.1) from either GenBank[4] or RefSeq[5], based on GTDB assembly entries. This resulted in a dataset of 24,402 species with their corresponding representative genomes.

##### Bioinformatically estimated growth rate data compilation

We compiled bioinformatically estimated maximum growth rates (i.e., minimum doubling time) for the previously assembled 24,402 species from the *Phydon* database[6]. *Phydon* is a genome-based framework that integrates codon usage patterns with phylogenetic information to predict maximum growth rates of microbes, providing temperature-corrected estimates for 111,349 microbial species. For comparison, we also considered *gRodon*, a widely used genome-based approach that estimates growth rates solely from codon usage patterns[7]. Using the experimentally measured growth rates compiled earlier as a benchmark, we found that *Phydon* outperformed *gRodon* in accurately predicting species' growth rates (Figure S11). Accordingly, we used Phydon-derived growth rate estimates for subsequent analyses.

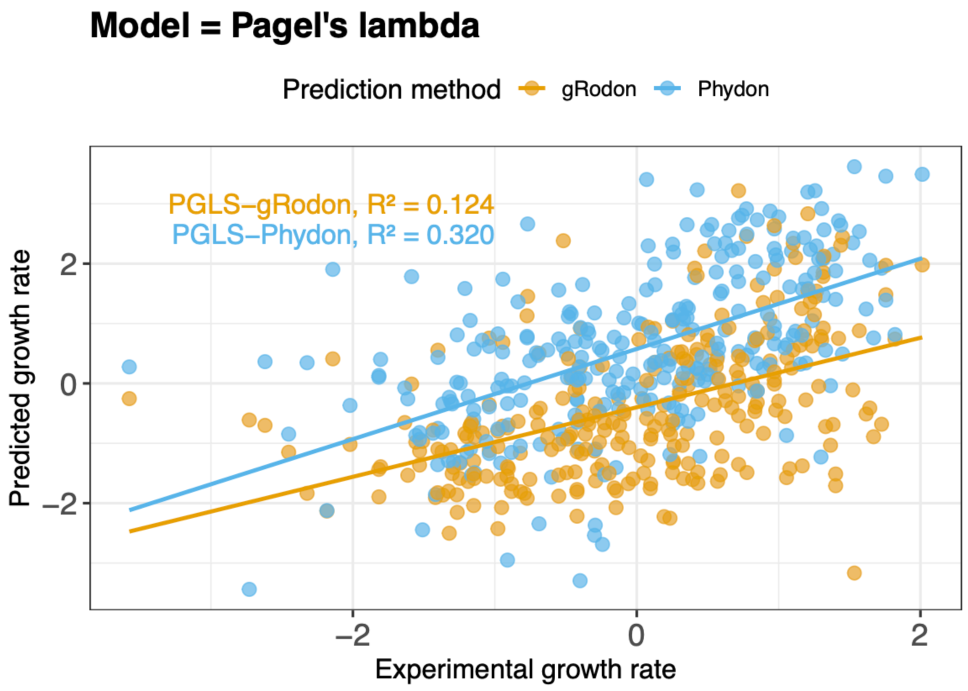

**Figure S11.** The performance of two bioinformatic tools, *Phydon* and *gRodon*, for inferring microbial growth rate was evaluated using experimentally measured growth rates as a benchmark. Correlations between predicted and observed growth rates across species (n = 283) were quantified using phylogenetic generalized least squares (PGLS) regression. Both the PGLS coefficients and $R^2$ values were compared between the two methods, with *Phydon* exhibiting superior predictive accuracy relative to *gRodon*.

**Trait definition and data processing**

For the supplementary analysis, data on the 11 associated traits for 24,402 species were defined, obtained, or calculated using the same procedures applied to species with experimentally measured growth rates (see Methods). This ensures consistency in trait definitions and data processing across datasets. The phylogenetic distribution of bioinformatically estimated growth rates, along with the 11 associated traits, is shown in Figure S12.

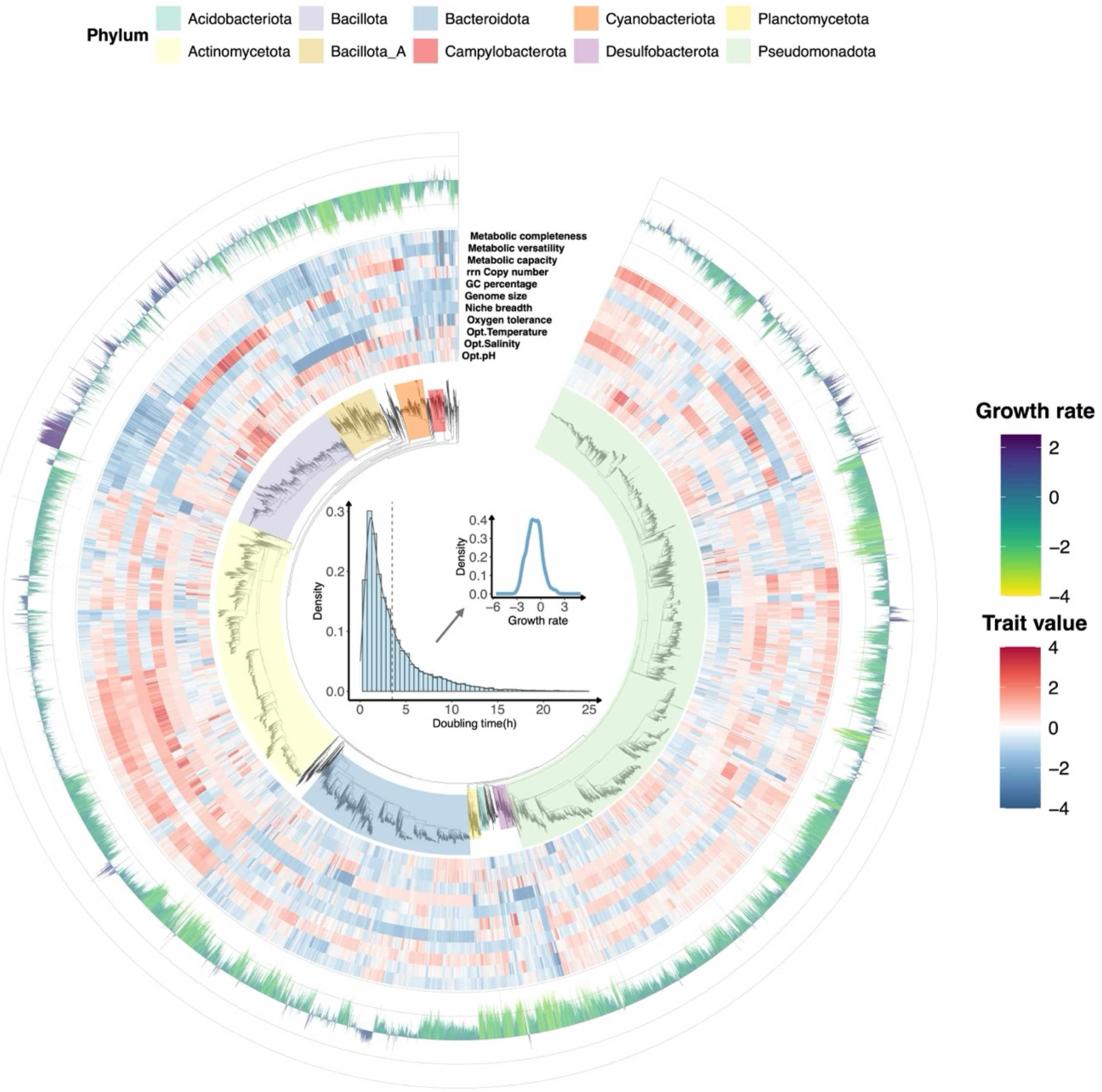

**Figure S12.** Phylogenetic distribution of bioinformatically estimated growth rates and associated traits across 24,402 species. The 11 associated traits were obtained or calculated consistently with the procedures used for species with experimentally measured growth rates. Species' minimum doubling times were converted to growth rates using a negative logarithm transformation.

**Mode of evolution**

Using the data compiled in the previous steps, we modelled the evolution of each trait independently and compared three alternative models of trait evolution — Brownian motion (BM), Ornstein–Uhlenbeck (OU), and early burst (EB) — to characterize the evolutionary mode of bioinformatically estimated growth rate and the 11 associated traits across 24,402 species.

We found that bioinformatically estimated growth rate and 9 of the associated traits were best described by the OU model, whereas two traits, GC percentage and oxygen tolerance, were best described by the EB model. These results suggest that growth rate and most associated traits evolved under stabilizing evolutionary processes, broadly consistent with the pattern observed in the main analysis based on experimentally measured growth rates (Figure S13).

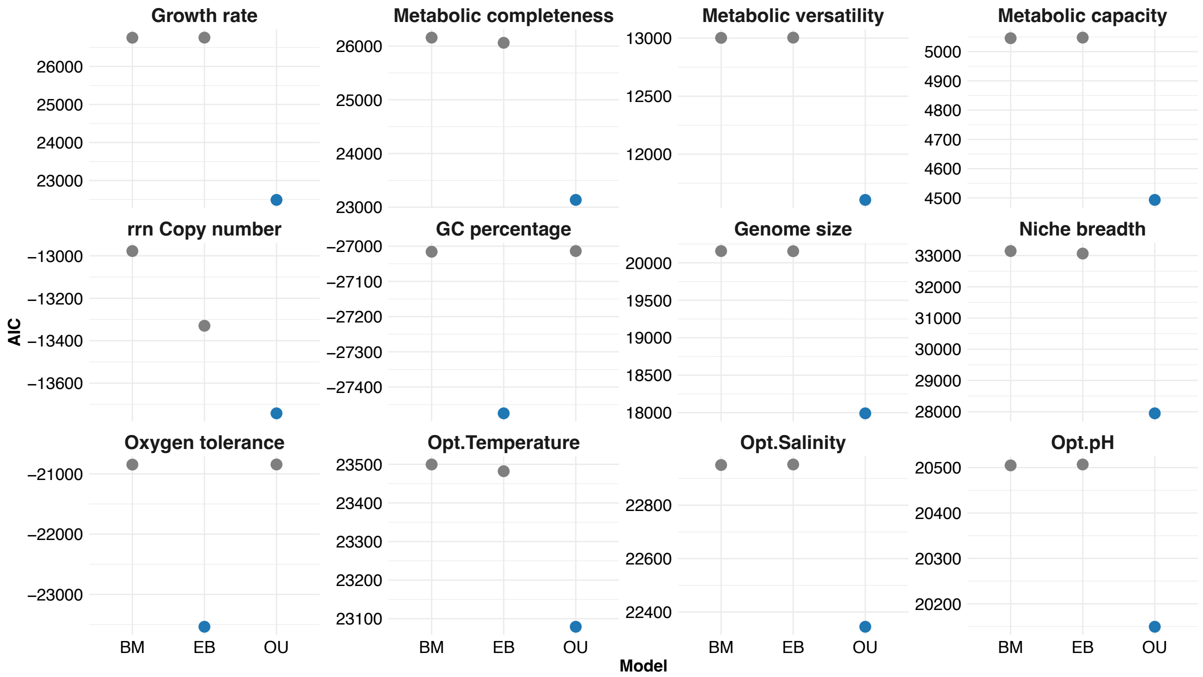

**Figure S13.** Comparison of goodness-of-fit for each trait among three commonly used trait evolution models (BM, EB, OU) based on a larger dataset with 24,402 species. AIC values are plotted for each model across the traits, with lower AIC values indicating a better model fit. The blue dot represents the model that best explains each trait.

For traits whose evolutionary mode was best described by an OU process, we further compared their evolutionary half-lives ($t_{1/2}$) to evaluate the relative tempo of evolution across traits. In this dataset, growth rate had the second shortest half-life, corresponding to 7.996% of the total tree height, whereas niche breadth had the shortest half-life, corresponding to 6.141% of the total tree height (Figure S14). This differs slightly from the main analysis, in which experimentally measured growth rate had the shortest half-life (Table 1). However, the two analyses are not contradictory. In both datasets, growth rate showed a relatively short evolutionary half-life compared with most other traits, indicating rapid return toward its evolutionary optimum. The difference in rank may reflect differences in taxonomic sampling, dataset size, and the additional uncertainty associated with bioinformatically estimated growth rates.

| | | Model Parameters | | | | | OU Dynamics | |
|---|---|---|---|---|---|---|---|---|
| Predictor | Best model | σ² (BM) | σ² (OU) | α (OU) | b (EB) | σ² (EB) | t½ (OU) | t½ (% tree height) |
| Growth rate | **OU** | | 8.336 | 8.669 | | | 0.080 | 7.996 |
| Metabolic completeness | **OU** | | 3.158 | 2.414 | | | 0.287 | 28.708 |
| Metabolic versatility | **OU** | | 4.427 | 3.555 | | | 0.195 | 19.496 |
| Metabolic capacity | **OU** | | 7.999 | 6.715 | | | 0.103 | 10.322 |
| rrn Copy number | **OU** | | 0.784 | 0.000 | | | 4,713,769.512 | 471,376,951.208 |
| GC percentage | **EB** | | | | −0.160 | 6.014 | | |
| Genome size | **OU** | | 1.521 | 2.815 | | | 0.246 | 24.628 |
| Niche breadth | **OU** | | 11.343 | 11.287 | | | 0.061 | 6.141 |
| Oxygen tolerance | **EB** | | | | −3.338 | 8.118 | | |
| Opt.Temperature | **OU** | | 6.694 | 2.160 | | | 0.321 | 32.083 |
| Opt.Salinity | **OU** | | 6.813 | 3.331 | | | 0.208 | 20.807 |
| Opt.pH | **OU** | | 6.011 | 2.464 | | | 0.281 | 28.136 |

**Figure S14.** Trait-evolution model fitting for each trait based on a larger dataset with 24,402 species. The best model for each trait is identified as either BM, OU, or EB, with the corresponding model parameters displayed for each evolutionary model. Parameter interpretations — σ²(BM), σ²(OU), σ²(EB): the variance in the evolutionary process for each model, describing how much the trait is expected to change over time; *b*(EB): the decay parameter of EB model, determining how quickly the rate of evolution decreases over time; α(OU): the rate of adaption of a trait under the OU process; $t_{1/2}$(OU): The half-life ($t_{1/2}$) of OU models, representing the time required for a trait to evolve halfway to its optimal state. The $t_{1/2}$ is calculated using the formula: $t_{1/2} = \frac{ln2}{\alpha}$, then is expressed as a fraction of total tree height.

We further validated the consistency of evolutionary rates estimated from OU models fitted to experimentally measured and bioinformatically estimated trait datasets. For traits whose evolutionary dynamics were best described by an OU process, the estimated OU half-lives showed a strong correlation between the two fitted models, indicating that the inferred evolutionary tempo was robust to the source of trait data (Figure S15).

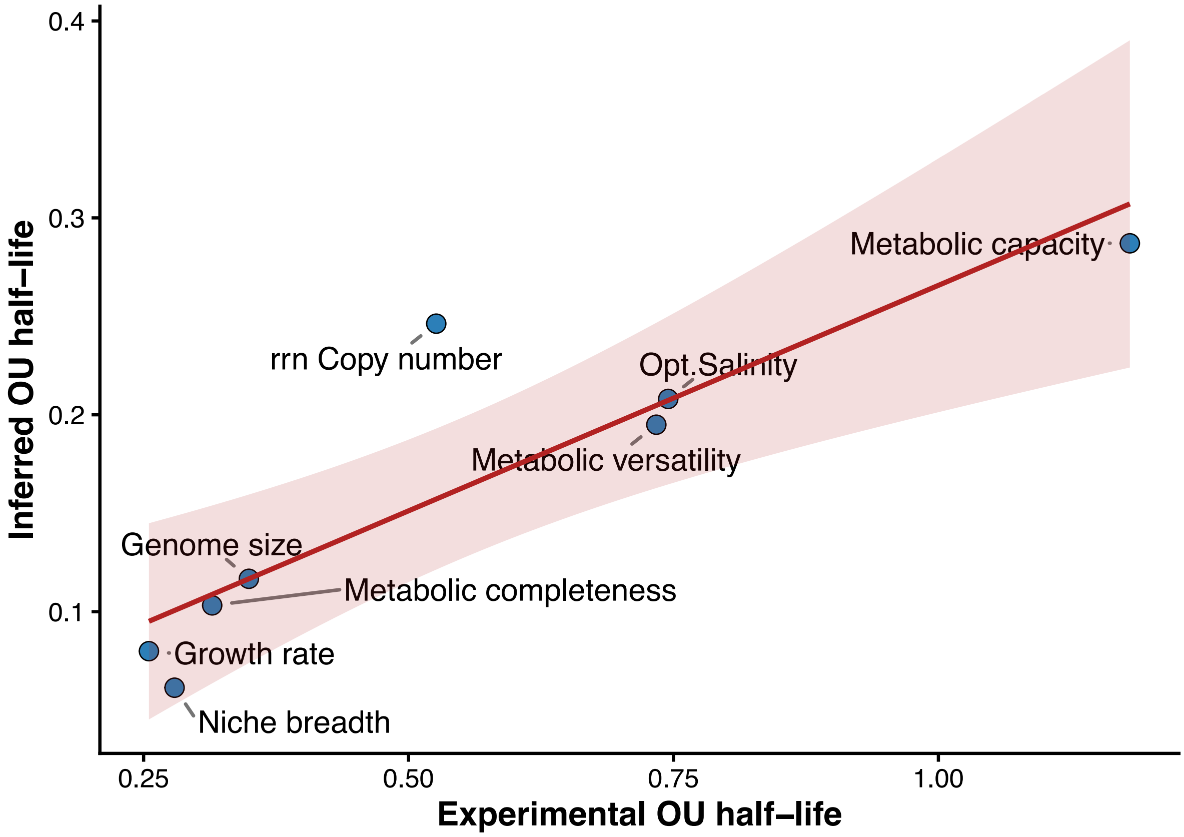

**Figure S15.** Scatter plot showing the relationship between trait evolutionary half-lives estimated using experimentally compiled data (x-axis) and bioinformatically inferred data (y-axis) for traits whose evolutionary dynamics were best described by an OU model. Each point represents one trait. The red line represents the linear regression between the two estimates, and the shaded region indicates the 95% confidence interval. The strong positive correlation demonstrates the consistency of evolutionary tempo inferred from experimentally measured and computationally estimated trait values.

#### Supplementary analysis 2: PGLS analyses of correlations between growth rate and associated traits

To compare with the patterns obtained using the multivariate Ornstein–Uhlenbeck (mvOU) framework, we applied phylogenetic generalized least squares (PGLS) analyses to evaluate the association between growth rate and each of the 11 associated traits while accounting for phylogenetic non-independence among species[8]. All PGLS analyses were performed under a Pagel's λ phylogenetic covariance structure using the R package "*phylolm*" (version 2.6.5)[9].

**PGLS-based associations**

Using experimentally measured growth-rate data, we found that metabolic completeness, *rrn* copy number, and optimum temperature were significantly positively correlated with growth rate across 283 species. The remaining traits showed no significant correlations with growth rate in this dataset (Figure S16). These three positive associations were also recovered when using bioinformatically estimated growth rates across 24,402 species (Figure S16). In addition, analyses based on the larger bioinformatically estimated dataset identified significant positive correlations between growth rate and metabolic versatility, GC percentage, genome size, niche breadth, and optimum salinity (Figure S16). The associations recovered in both datasets provide support for the robustness of key correlations, whereas the additional correlations detected only in the bioinformatically estimated dataset may reflect increased statistical power from broader taxonomic sampling.

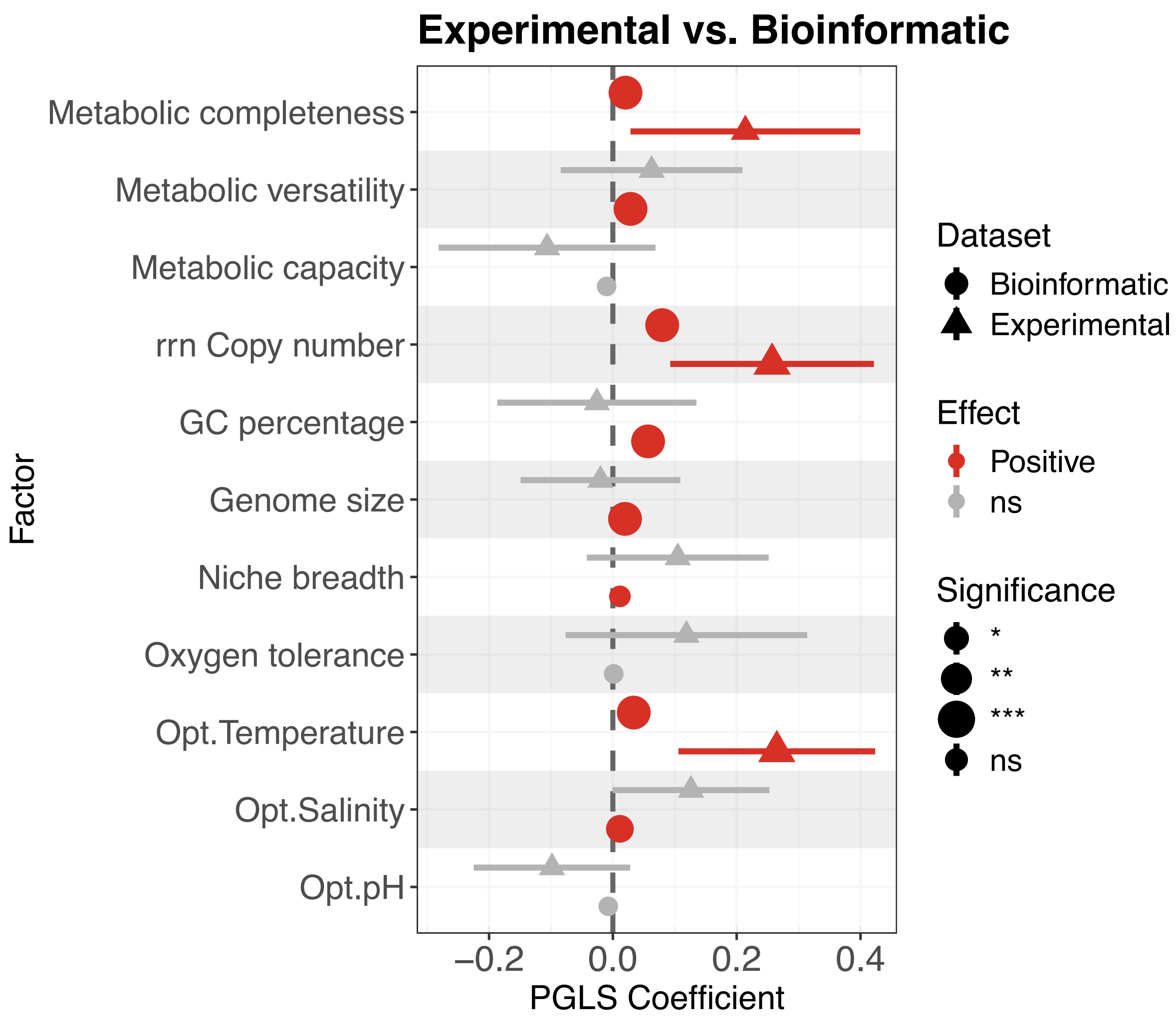

**Figure S16.** Comparison of PGLS results for experimentally measured versus bioinformatically inferred growth rates. PGLS models were fitted with growth rate as the response variable using experimentally measured data (n = 283; triangles) and bioinformatically inferred data (n = 24,402; circles). Each point represents the estimated regression coefficient for the association between a trait and growth rate, with 95% confidence intervals shown.

**Simulation on the relationships between mvOU and PGLS**

To better understand how parameter structure in a mvOU model can reflect the evolutionary association estimated by PGLS, we conducted a simulation on a fixed phylogenetic tree. The purpose of this simulation was to evaluate whether a pairwise trait association, as estimated by PGLS, can reflect different coevolutionary scenarios implied by different underlying bivariate OU processes.

We simulated two continuous traits, denoted Trait 1 and Trait 2, on a randomly generated phylogeny with 100 tips. Trait evolution was modelled using a bivariate OU process with a single global adaptive regime. The root state and optimum were both fixed at zero for the two traits, so that any association between simulated trait values arose from the covariance structure generated by the OU dynamics rather than from differences among selective regimes or initial trait means.

The diagonal elements were held constant, meaning that the intrinsic strength of attraction toward the optimum was fixed for both traits. The diffusion matrix ($\Sigma_{yy}$ matrix) was also fixed as diagonal, with no correlated diffusion for the two traits. Therefore, the simulation isolated the effect of the off-diagonal elements of the adaptation matrix of a given bivariate OU model. These off-diagonal elements represent cross-trait coevolutionary coupling: $\boldsymbol{A}_{12}$ describes how deviations in Trait 2 contribute to the evolution of Trait 1, whereas $\boldsymbol{A}_{21}$ describes how deviations in Trait 1 contribute to the evolution of Trait 2. We explored a grid of values for $\boldsymbol{A}_{12}$ and $\boldsymbol{A}_{21}$, ranging from $-1$ to 1 in increments of 0.02. For each combination of off-diagonal parameters, we simulated replicate bivariate trait datasets (n = 10) under the corresponding OU model. Parameter combinations that produced an unstable OU process were excluded using an eigenvalue-based stability check, requiring all eigenvalues of the drift matrix to have positive real parts. For each successfully simulated dataset, we fitted a PGLS model to estimate the association of the two traits under a Brownian-motion phylogenetic covariance structure, the estimated PGLS regression coefficient was recorded. We then summarized the mean PGLS slope across replicates for each combination of $\boldsymbol{A}_{12}$ and $\boldsymbol{A}_{21}$.

The simulation showed that similar PGLS coefficients could arise from different combinations of $\boldsymbol{A}_{12}$ and $\boldsymbol{A}_{21}$ (Figure S17). In the heat map, broad regions of the OU parameter space produced comparable positive or negative PGLS slopes, rather than each slope value corresponding to a single unique combination of cross-trait parameters. This demonstrates that a PGLS coefficient summarizes the marginal association between observed tip values, but does not by itself identify the specific evolutionary dynamics responsible for that association. Therefore, a positive or negative PGLS slope may be generated by particular combinations of OU parameters, and different OU parameter combinations may produce similar marginal regression patterns (Figure S17).

To summarize, our simulation showed that PGLS and multivariate OU models answer related but distinct questions: PGLS can reveal whether an association is present, but multivariate OU modelling is needed to evaluate the evolutionary dynamics that may have generated that association.

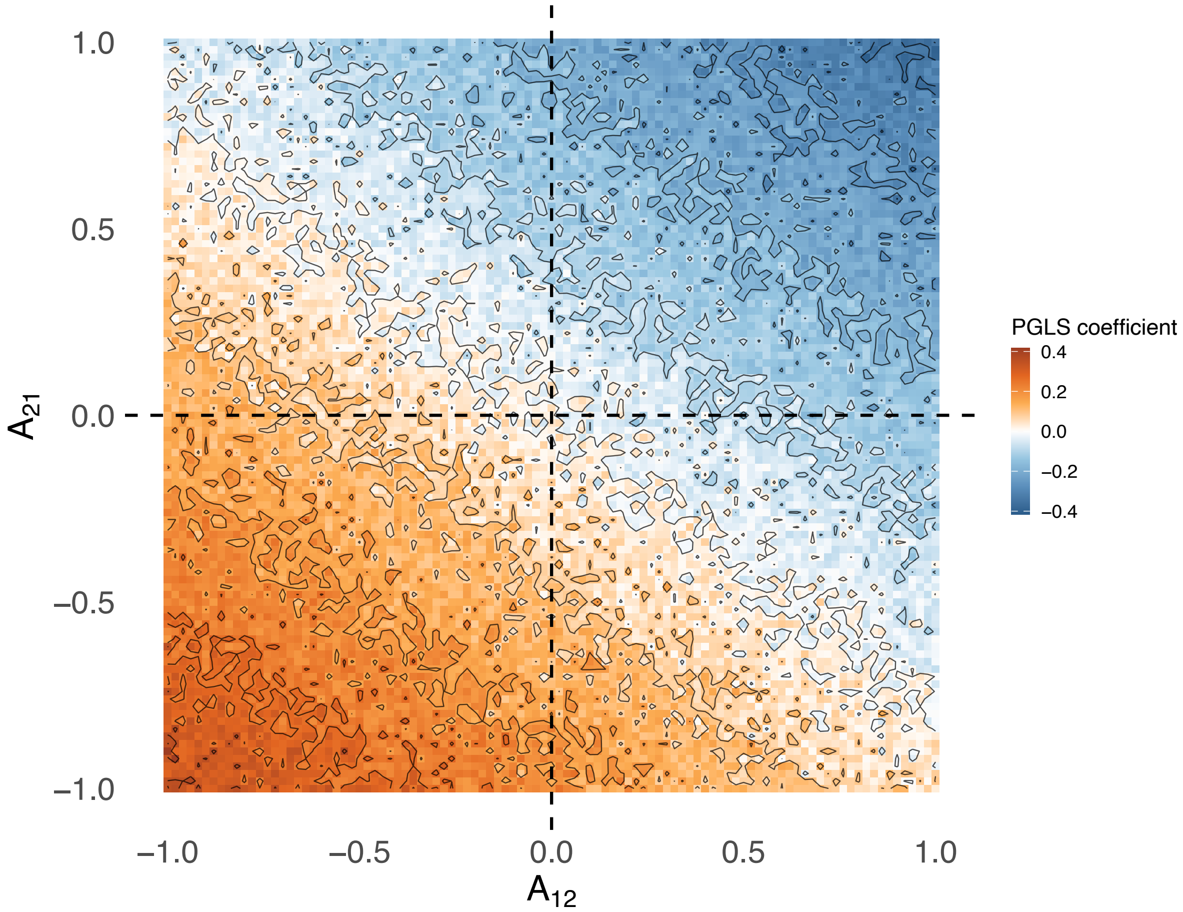

**Figure S17.** Relationship between bivariate OU dynamics and PGLS-estimated trait association. Heat map showing the mean PGLS coefficient for the regression of Trait 2 on Trait 1 across simulated datasets generated from bivariate OU processes with varying off-diagonal adaptation matrix parameters, $\boldsymbol{A}_{12}$ and $\boldsymbol{A}_{21}$. For each parameter combination, trait values were simulated on a fixed phylogenetic tree and analysed using PGLS under a Brownian-motion covariance structure. Colours indicate the mean PGLS coefficient, with orange representing positive associations and blue representing negative associations.

#### Comparison of pairwise PGLS coefficients with the stationary covariance matrices of the mvOU model

Under the multivariate OU framework, the stationary covariance matrix of a fitted model describes the variance–covariance structure that the trait vector is expected to attain once the process has reached evolutionary equilibrium[10]. Its diagonal entries are the stationary variances of the individual traits, and its off-diagonal entries are the stationary covariances between trait pairs; the sign of an off-diagonal entry indicates whether the two traits are inferred to vary in the same direction or in opposite directions at the equilibrium under the fitted mvOU model[10].

For each growth rate–trait pair we therefore compared two quantities that describe the same association from different perspectives: the pairwise PGLS coefficient for that pair, and the corresponding off-diagonal element of the mvOU-derived stationary covariance matrix (Figure S18). The two differ in their theoretical underpinnings and in what they estimate. Nevertheless, a qualitative comparison of their signs is meaningful because both provide time-independent summaries of the direction of pairwise trait association: the PGLS coefficient describes the phylogenetically adjusted association among observed species, whereas the stationary covariance describes the model-implied association between the same traits at evolutionary equilibrium. PGLS returns a phylogenetic regression slope, a phenomenological estimate of the association observed across species after accounting for shared ancestry. The stationary covariance, by contrast, is a model-implied expectation of the long-run covariance between the two traits at evolutionary equilibrium under the fitted OU process.

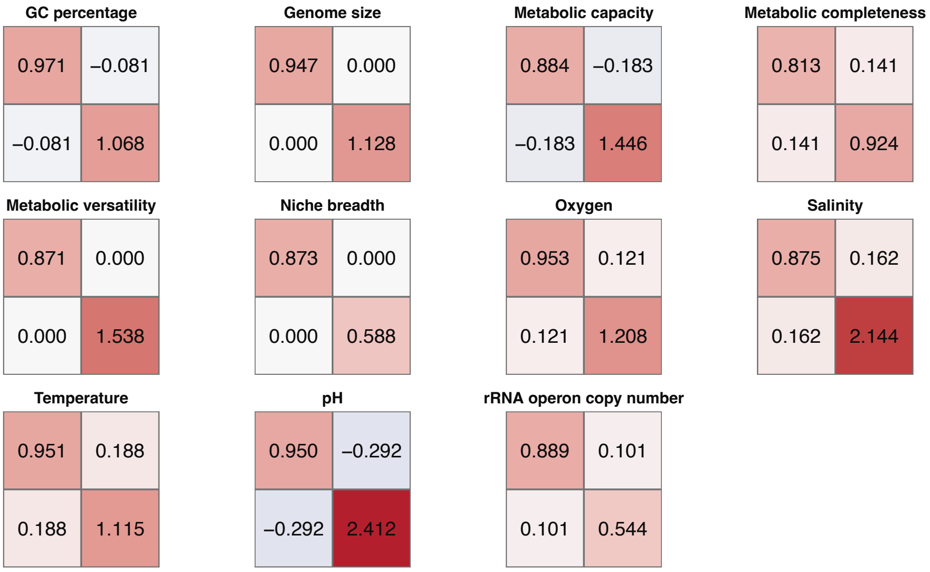

**Figure S18.** Stationary covariance matrices of the best-fitting mvOU model for each of the growth rate–trait pairs. Each panel shows the 2 × 2 stationary covariance matrices for one pair, with growth rate given first in both the rows and the columns and the trait of interest given second. Diagonal entries are stationary variances and off-diagonal entries are stationary covariances; a positive covariance indicates that the two traits are inferred to vary in the same direction at evolutionary equilibrium and a negative covariance that they vary in opposite directions.

Because the two quantities are expressed on different scales and are not estimates of a common parameter, they cannot be compared quantitatively; we therefore compared them qualitatively, on the basis of the sign of the association and of whether each was statistically supported. To place the stationary covariances on a scale that is directly comparable among trait pairs, we converted each fitted variance–covariance matrix into a correlation matrix with the R function "cov2cor". In the resulting matrix the diagonal entries equal 1 and the off-diagonal entries lie between −1 and 1 and express the strength as well as the direction of the model-implied stationary association.

We then compared, for each trait pair, the sign and statistical support of the PGLS coefficient with the sign of the corresponding off-diagonal element of the stationary correlation matrix derived from the corresponding best-support mvOU model (Figure S19). We found that the mvOU framework identified a non-zero stationary covariance for eight pairs, whereas pairwise PGLS found a significant association for three, so five pairs that carry an inferred evolutionary dependency under the fitted OU model are not detected by PGLS (Figure S19). For example, neither metabolic capacity or optimum pH shows a significant association with growth rate under PGLS, yet both have non-zero stationary covariances under the fitted mvOU model. This pattern indicates that pairwise PGLS is not by itself sufficient to capture the evolutionary dependencies between traits, because it does not model the coevolutionary relationship that, within the OU framework, determines whether and how strongly the traits are coupled over macroevolutionary time.

| Trait pair | PGLS | mvOU | Agreement |
|---|---|---|---|
| Growth rate ~ Opt.pH | ns | – | NA |
| Growth rate ~ Opt.Salinity | ns | + | NA |
| Growth rate ~ Opt.Temperature | + | + | Yes |
| Growth rate ~ Oxygen tolerance | ns | + | NA |
| Growth rate ~ Niche breadth | ns | 0 | NA |
| Growth rate ~ Genome size | ns | 0 | NA |
| Growth rate ~ GC percentage | ns | – | NA |
| Growth rate ~ rrn Copy number | + | + | Yes |
| Growth rate ~ Metabolic capacity | ns | – | NA |
| Growth rate ~ Metabolic versatility | ns | 0 | NA |
| Growth rate ~ Metabolic completeness | + | + | Yes |

**Figure S19.** Qualitative comparison of the pairwise PGLS coefficient and the stationary covariance of the mvOU model for each of the 11 growth–trait pairs. The PGLS column gives the sign of the PGLS coefficient for that pair, with "+" and "−" denoting a statistically significant positive or negative slope ($p < 0.05$) and "ns" denoting a slope that is not significant. The mvOU column gives the sign of the corresponding off-diagonal element of the model-implied stationary correlation matrix, with "+" and "−" denoting a positive or negative covariance and "0" denoting a covariance constrained to zero by the fitted model. The Agreement column reports "Yes" where the two signs concur and both quantities are informative, and "NA" where the comparison is uninformative because the PGLS coefficient is not significant or the stationary covariance is constrained to zero.

#### Supplementary analysis 3: uncovering the multivariate coevolution of traits.

Our main analyses evaluated the coevolutionary relationship between growth rate and each associated trait separately. However, these traits may also covary or coevolve with one another. In complex scenarios where multiple genomic, metabolic, ecological, and environmental traits jointly influence growth rate, bivariate coevolutionary models may oversimplify the underlying evolutionary dynamics[11]. To address this limitation, we applied a multivariate Ornstein–Uhlenbeck (mvOU) framework to model the joint coevolution of growth rate and multiple interacting traits. This approach allows traits to coevolve and potentially influence one another, and tests whether growth-rate evolution is shaped by the combined effects of genomic, metabolic, ecological, and environmental traits (Figure 1b).

**Generating composite trait axes**

Including all 11 associated traits together with growth rate in a single model would require modelling 12 variables simultaneously. Such a model would be highly complex, difficult to interpret, and associated with an impractically large number of candidate models describing alternative coevolutionary relationships among traits. In addition, many of these traits are likely to be correlated, further complicating model estimation and interpretation.

To reduce model complexity, improve interpretability, and account for covariance among correlated variables within each trait category, we performed phylogenetically informed principal component analyses (pPCA) using the standardized data of 283 species with experimental growth rate for variables assigned to each category (Figure 2c). Specifically, pPCA were conducted separately on (i) the three genomic characteristic traits (GC), (ii) the three metabolic potential traits (MP), and (iii) the four environmental tolerance traits (ET). Niche breadth, the only ecological variable, was retained as an independent predictor (Figure 1b). For each trait category, we extracted the first phylogenetic principal component (pPC1), which captured the primary axis of covariation, and used it as a composite variable in subsequent analyses (Figure S20). These analyses were carried out using the "phyl.pca" function, with correlation structure specified as "lambda", in the R package "*phytools*" (version 2.4.4[12]). These pPC1 scores captured a substantial proportion of variance (55.4%, 59.9%, and 40.2%, respectively) and provided biologically interpretable composite variables for testing multivariate coevolution with growth rate (Figure S20).

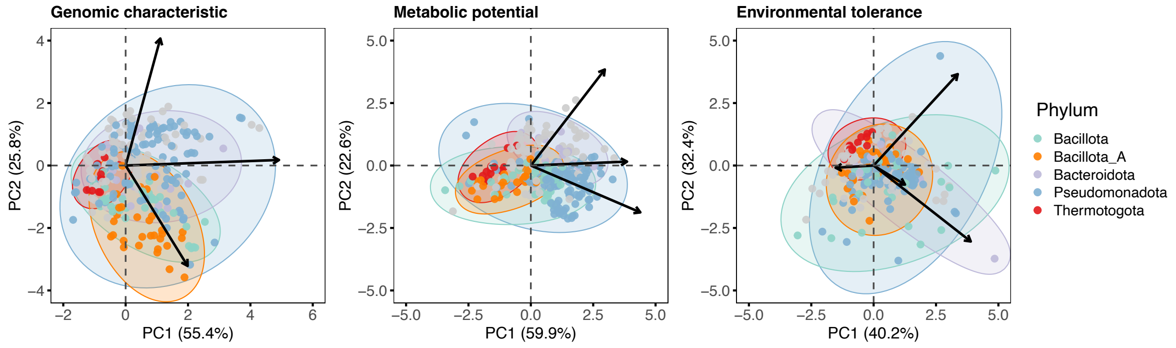

**Figure S20.** Multivariate coevolution between growth rate and major trait axes. **(a)** Phylogenetically informed PCA of genomic characteristics, metabolic potential, and environmental tolerance. For each trait category, the first principal component (pPC1) captured the dominant axis of covariation (55.4%, 59.9%, and 40.2% of the variance, respectively) and was used as a composite variable in subsequent mvOU analyses. Niche breadth was retained as an independent ecological predictor.

#### Hypotheses regarding direct and indirect coevolutionary relationships

We formulated 45 candidate models, each represented as a directed acyclic graph (DAG), to evaluate distinct hypotheses regarding the multivariate relationships among the five variables: growth rate (GR), genomic characteristic (GC), metabolic potential (MP), environmental tolerance (ET), and niche breadth (NB; Figure S21). Each model specifies a distinct hypothetical scenario, differing in which variables are assumed to exert direct effects on growth rate and which act as mediators transmitting the effects of other variables. For example, model_H01 represents a fully direct scenario in which all variables independently influence growth rate without mediation, whereas model_H02 posits that the effect of genomic characteristics on growth rate is indirect, and mediated by metabolic potential, while other variables retain direct effects (Figure S21).

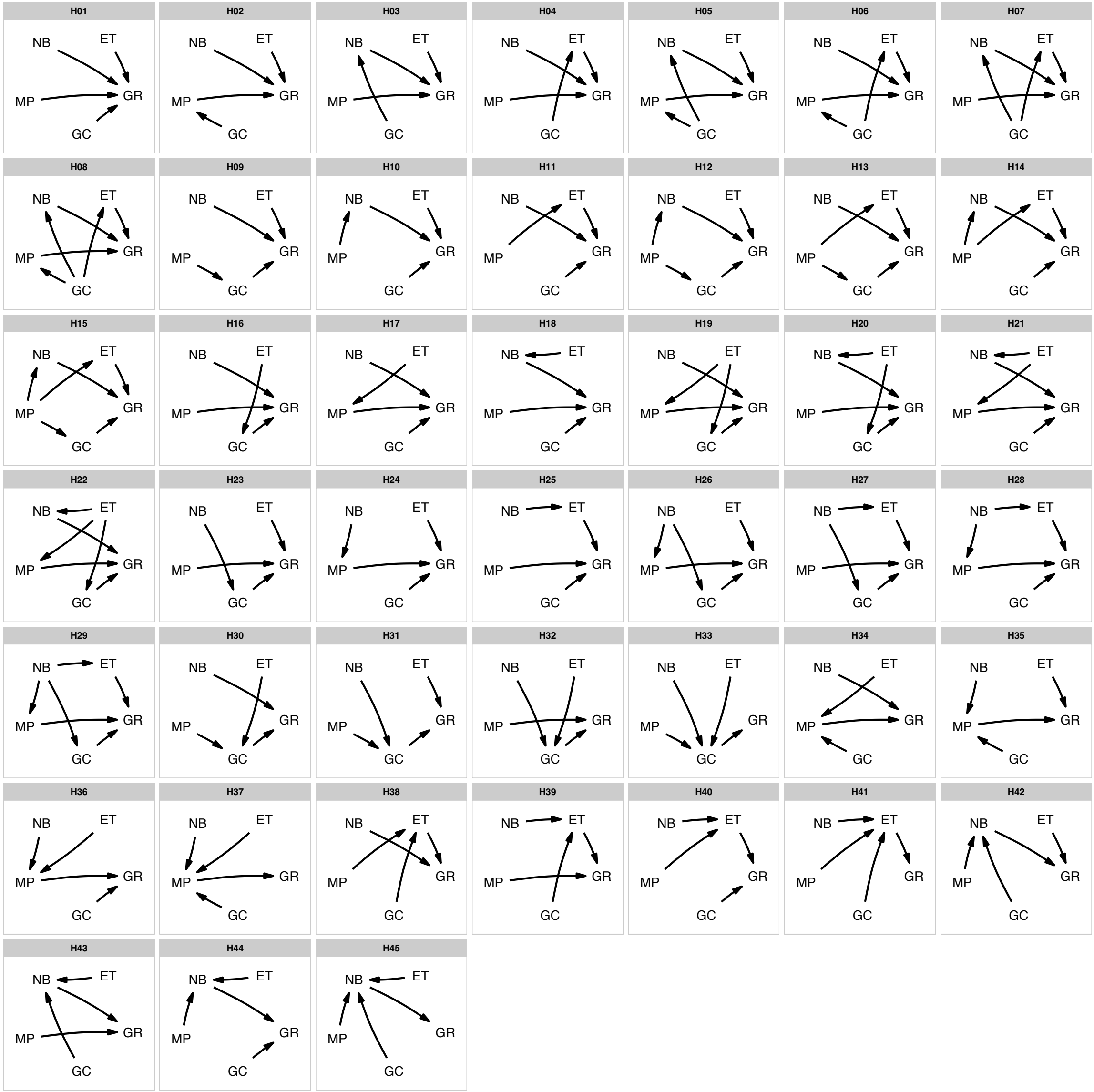

**Figure S21.** The 45 candidate models represent alternative hypotheses about the multivariate relationships among the five variables analysed in this study. Each model is illustrated as a directed acyclic graph (DAG), showing hypothesized direct and indirect relationships among variables. Variable abbreviations: GR = growth rate; GC = genomic characteristics; MP = metabolic potential; ET = environmental tolerance; NB = niche breadth.

#### Fitting mvOU models to infer direct and indirect coevolutionary relationships among traits

Our first step was to compare the relative likelihood of the 45 previously defined hypotheses under a mvOU framework (Figure S21), which represent hypotheses for the direct and indirect relationships among five variables. We implemented all model fitting using the R package "*mvSLOUCH*" (2.7.6[10]).

In mvSLOUCH, the structure of each candidate DAG is encoded in the $\boldsymbol{A}$ matrix through parameter constraints: (i) If a DAG specified no direct link between two nodes (traits), the corresponding off-diagonal entry in the $\boldsymbol{A}$ matrix was fixed at 0. (ii) If a DAG specified a direct link between two nodes, the corresponding off-diagonal entry in the $\boldsymbol{A}$ matrix was set to 'NA', allowing it to be freely estimated[10]. Model selection was based on the AICc.

For this initial model comparison step, our aim was to distinguish the direct and indirect associations among traits. Therefore, all diagonal entries of $\boldsymbol{A}$ were constrained to be positive, ensuring each trait approaches to its own optimum, while all non-zero off-diagonal entries were left as 'NA' (free to fit), allowing the data to determine the sign and strength of each direct coevolutionary coupling without prior assumption.

We also included two additional reference models. First, we fitted a simple null model in which all entries of the $\boldsymbol{A}$ matrix were fixed at 0, representing a scenario with no self-adaptation and no coevolution among traits. Second, we fitted a fully parameterized model in which all entries of the $\boldsymbol{A}$ matrix were freely estimated. If model selection based on information criteria favours the null model, this may indicate that the data contain insufficient information to support reliable parameter estimation. Conversely, if the fully parameterized model is favoured, this may suggest that the best-supported evolutionary scenario is not captured by the predefined set of candidate hypotheses[13].

To ensure robust parameter estimates and confident model selection, each model configuration was run 100 times with different starting points, obtained from preliminary fits with $\boldsymbol{A}$ set to "*DecomposablePositive*".

We tested whether the traits evolved with correlated or independent stochastic perturbations by fitting each model with two parameterizations of the $\Sigma_{yy}$ diffusion matrix: 'diagonal' (independent perturbations) and 'upper triangular' (correlated perturbations). Across all candidate models, the 'diagonal' parameterization was consistently favoured by smaller AICc, indicating that stochastic perturbations act independently on each trait (Figure S22). Consequently, we only analysed models with 'diagonal' $\Sigma_{yy}$ matrix.

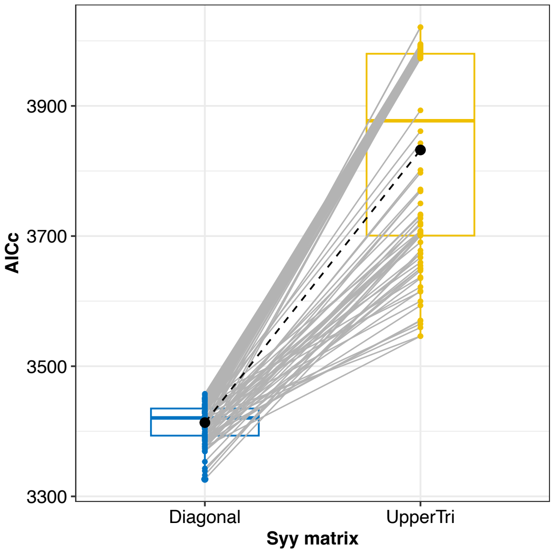

**Figure S22.** Model selection for the mode of stochastic perturbations across 47 models (45 candidate models + 2 reference models). Dots represent the AICc scores for models with either a diagonal or upper triangular $\Sigma_{yy}$ matrix (the stochastic perturbation matrix). Models with diagonal $\Sigma_{yy}$ consistently yielded lower AICc values, indicating that these traits evolve primarily under independent stochastic perturbations rather than correlated selection.

For model selection, we also compared the goodness of fit between two selective-regime assumptions: an OU1 model, in which each trait evolves toward a single global optimum, and an OUM model, in which each trait evolves toward one of two optima defined by growth-rate regimes. The best-supported model was identified as the model with the lowest AICc value across all runs and candidate models.

The best-supported model was model H08 under the OU1 assumption, indicating that adding growth-rate-defined selective regimes did not improve model fit (Figure S23). We therefore retained this model for further parameterization in the subsequent step. We note that the joint evolution of the five traits could, in principle, be shaped not only by selective regimes defined by growth rate, but also by regimes associated with other traits or by interactions among multiple trait-defined regimes. Here, we considered only growth-rate-defined regimes to keep the analysis tractable. More complex selective-regime structures may provide additional explanatory power, but were beyond the scope of this analysis.

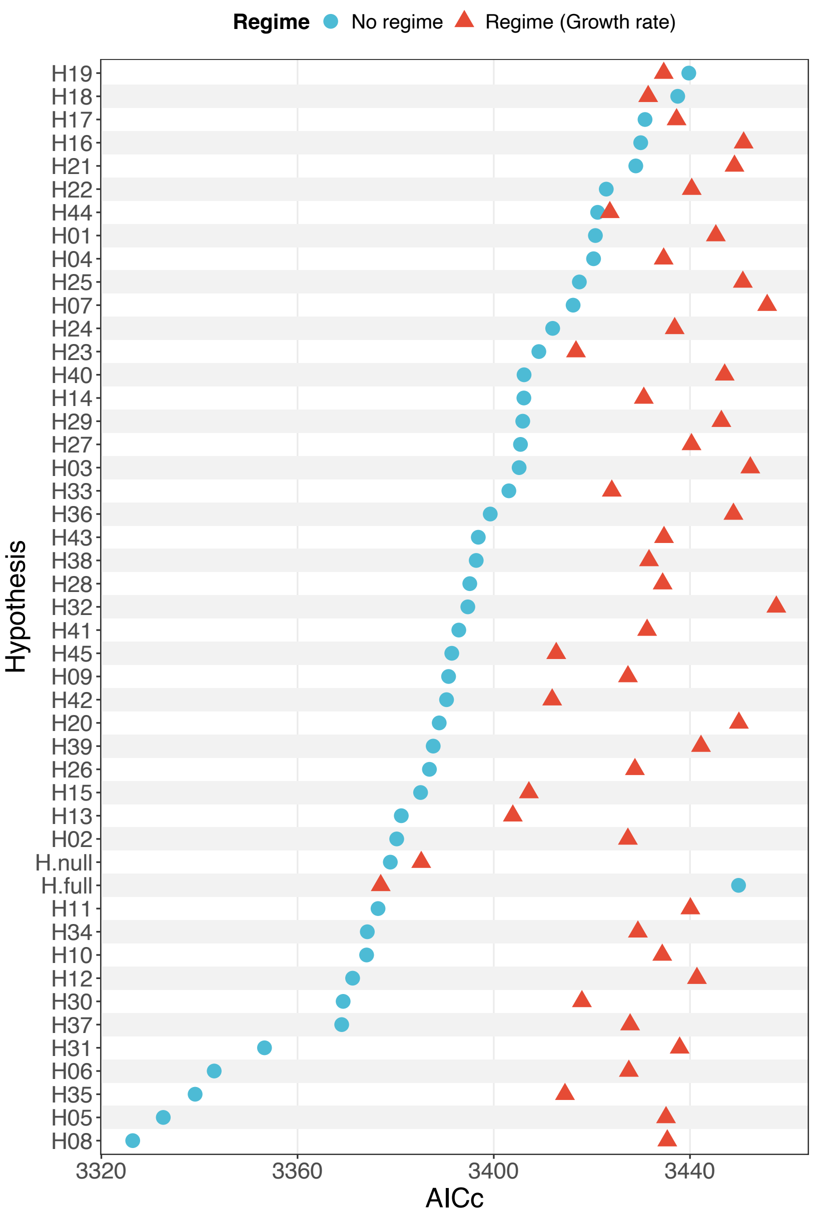

**Figure S23.** Model selection was performed across 47 models, comprising 45 candidate models and two reference models, under both OU1 and OUM assumptions. The OU1 assumption specifies no additional selective regimes, whereas the OUM assumption allows separate optima defined by growth-rate regimes. The best-supported model was model H08 under the OU1 assumption.

#### Identifying the direction of coevolution in the best-supported model

A subsequent step was performed to infer the most likely direction of coevolutionary coupling within the best-supported model (model H08) identified in the previous step. The asymmetry of the $\boldsymbol{A}$ matrix ($\boldsymbol{A}_{ij} \neq \boldsymbol{A}_{ji}$), provides a framework for this type of causal inference, as each possible direction of an arrow (e.g., Trait A → Trait B vs. Trait B → Trait A) represents a distinct biological hypothesis about which trait exerts evolutionary influence on the other[14]. To evaluate these alternatives, we treated the previously selected best DAG as a skeleton, and systematically tested all possible combinations of directions for its arrows, with each combination defining a distinct causal model. Because model H08 contains six direct coevolutionary links, and each link can be assigned one of two possible directions, this resulted in 64 candidate directional models (Figure S24).

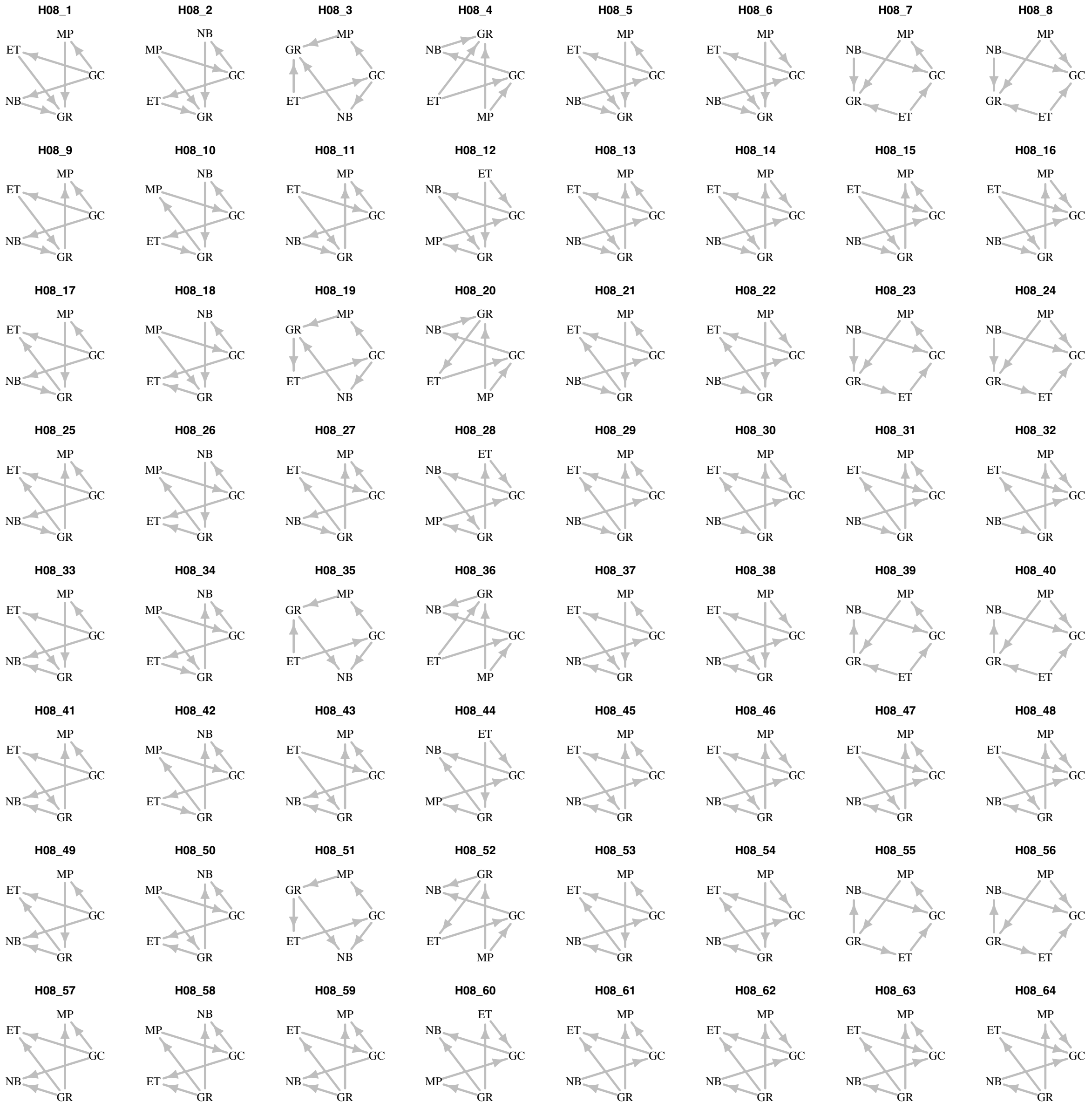

**Figure S24.** Candidate directional models derived from model H08.The 64 candidate directional models derived from the H08 model skeleton were evaluated to infer the most likely direction of coevolutionary coupling among traits. Models are labelled from H08_1 to H08_64, with each model representing a unique combination of directions for the six direct coevolutionary links in H08. Trait abbreviations: GR, growth rate; GC, genomic characteristics; MP, metabolic potential; ET, environmental tolerance; NB, niche breadth.

We compared these models using AICc to identify the directional structure best supported by the data. For each candidate directional model, we also compared two selective-regime assumptions: OU1 model vs. OUM model, in which trait optima are defined by growth-rate regimes. To ensure robust parameter estimates for this set of candidate models, each was run 100 times from different starting points, and the run with the lowest AICc was reported as the final best model. Model H08_60 under the OU1 assumption was selected as the best model, our conclusions were therefore drawn by analysing its $\boldsymbol{A}$ matrix, and the eigenvectors and eigenvalues of the $\boldsymbol{A}$ matrix from the selected final best model (Figure S25).

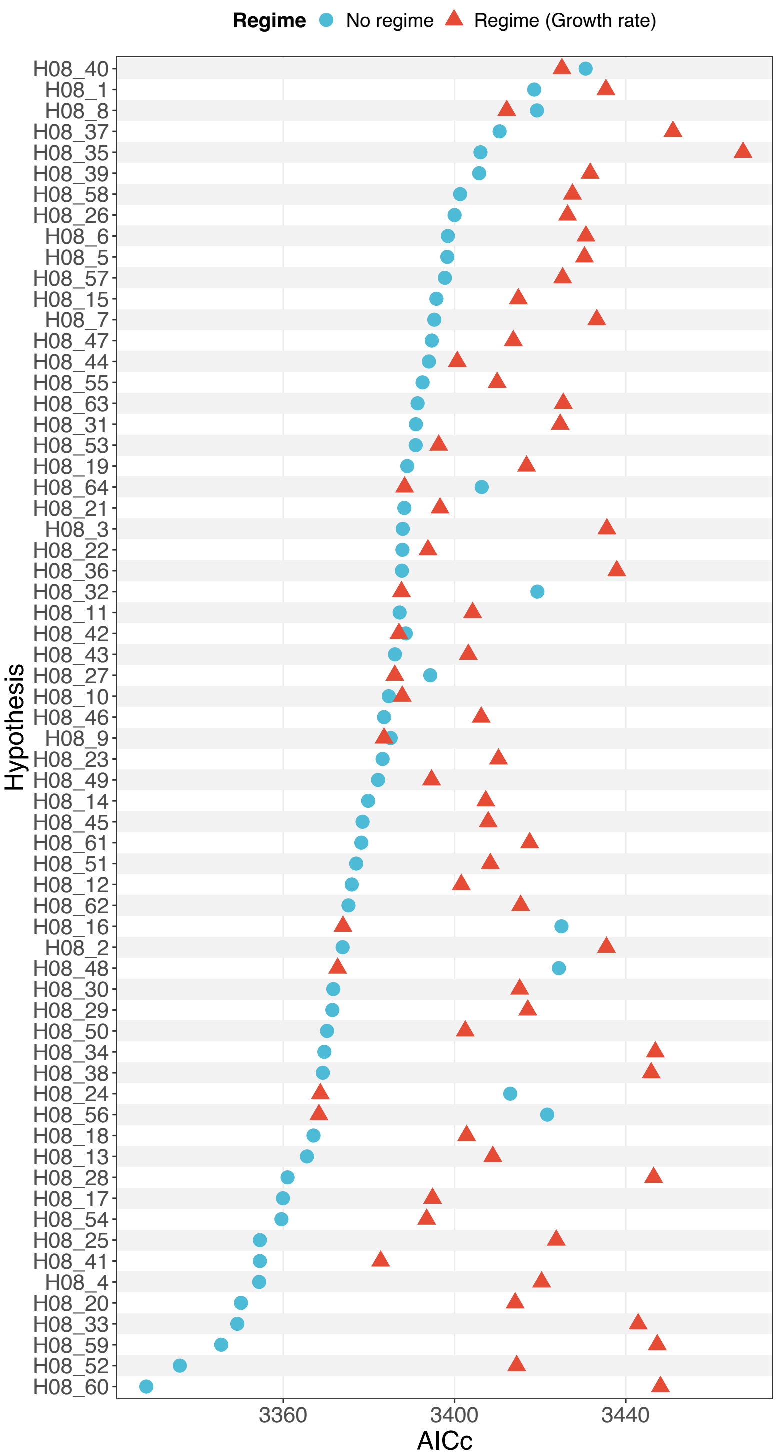

**Figure S25.** Model selection was performed across 64 directional models under both OU1 and OUM assumptions. The best-supported model was model H08_60 under the OU1 assumption.

#### Growth rate dominates the fastest-evolving mode

Analysis of the ***A*** matrix from the best-supported model (model H08_60) indicated that growth rate directly drives the evolution of metabolic potential, environmental tolerance, and niche breadth, whereas no direct coevolutionary relationship was identified between genomic characteristics and growth rate over instantaneous evolutionary time scales (Figure S26a). Instead, the evolution of genomic characteristics was influenced by metabolic potential and environmental tolerance, and in turn, genomic characteristics shaped the evolution of niche breadth (Figure S26a).

Eigen-decomposition of the fitted $\boldsymbol{A}$ matrix for the best-supported model showed that growth rate had the largest loading on the fastest-evolving mode ($\overrightarrow{e_1}$) over longer evolutionary timescales. Metabolic potential, environmental tolerance and niche breadth also loaded on $\overrightarrow{e_1}$, but in the opposite direction to growth rate (Figure S26b). By contrast, genomic characteristics loaded on $\overrightarrow{e_1}$ in the same direction as growth rate, indicating that although no direct coevolution is detected between them on instantaneous timescales, their evolutionary trajectories converge over longer timescales (Figure S26b).

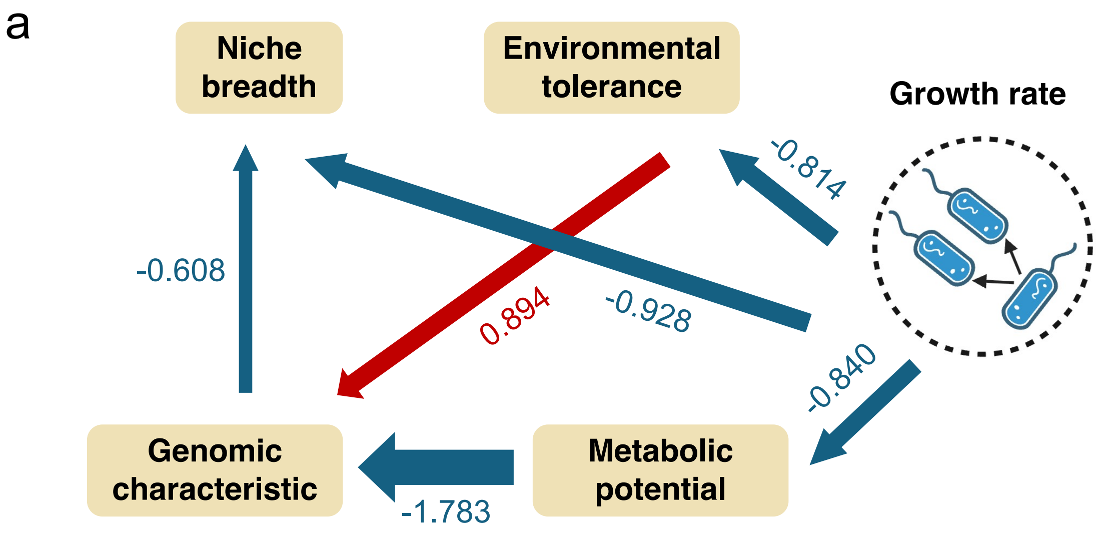

b

| Trait / Metric | $\vec{e1}$ | $\vec{e2}$ | $\vec{e3}$ | $\vec{e4}$ | $\vec{e5}$ |
|---|---|---|---|---|---|
| **Genomic characteristic** | 0.226 | -0.214 | 0.000 | 0.806 | -0.845 |
| **Metabolic potential** | -0.409 | 0.115 | 0.000 | 0.000 | 0.000 |
| **Environmental tolerance** | -0.212 | 0.000 | 0.000 | 0.000 | 0.272 |
| **Niche breadth** | -0.501 | 0.970 | **1.000** | 0.592 | -0.460 |
| **Growth rate** | 0.697 | 0.000 | 0.000 | 0.000 | 0.000 |
| **Eigenvalue ($\lambda$)** | 3.050 | 1.619 | 1.485 | 0.657 | 0.370 |
| **Half-life ($t_{1/2} = \frac{ln2}{\lambda}$)** | 0.227 | 0.428 | 0.467 | 1.055 | 1.875 |
| **Tree Height (%)** | 22.724 | 42.809 | 46.665 | 105.453 | 187.531 |

**Figure S26.** Multivariate coevolution between growth rate and major trait axes. (a) Multivariate mvOU coevolutionary structure inferred from the best-supported model. Directed arrows denote significant instantaneous coevolutionary effects among composite variables after jointly modelling all predictors. (b) Eigen-decomposition of the A matrix of the best model. Each eigenvector represents an independent adaptive mode in multivariate trait space, with its corresponding eigenvalue (λ) determining the rate at which that mode evolves towards the optimum. This rate is then summarized by the evolutionary half-life (ln2/λ), and its proportion of the total tree height. Trait loadings on each eigenvector are displayed. For each eigenvector, the trait that completely dominates the adaptive mode is highlighted in bold.

### References

1. Louca, S., Mazel, F., Doebeli, M. & Parfrey, L. W. A census-based estimate of Earth's bacterial and archaeal diversity. *PLOS Biology* **17**, e3000106 (2019).

2. Parks, D. H. *et al.* GTDB: an ongoing census of bacterial and archaeal diversity through a phylogenetically consistent, rank normalized and complete genome-based taxonomy. *Nucleic Acids Research* **50**, D785–D794 (2022).
3. Bowers, R. M. *et al.* Minimum information about a single amplified genome (MISAG) and a metagenome-assembled genome (MIMAG) of bacteria and archaea. *Nat Biotechnol* **35**, 725–731 (2017).
4. Benson, D. A. *et al.* GenBank. *Nucleic Acids Research* **41**, D36–D42 (2013).
5. O'Leary, N. A. *et al.* Reference sequence (RefSeq) database at NCBI: current status, taxonomic expansion, and functional annotation. *Nucleic Acids Research* **44**, D733–D745 (2016).
6. Xu, L., Zakem, E. & Weissman, J. L. Improved maximum growth rate prediction from microbial genomes by integrating phylogenetic information. *Nat Commun* **16**, 4256 (2025).
7. Weissman, J. L., Hou, S. & Fuhrman, J. A. Estimating maximal microbial growth rates from cultures, metagenomes, and single cells via codon usage patterns. *Proceedings of the National Academy of Sciences* **118**, e2016810118 (2021).
8. Cornwell, W. & Nakagawa, S. Phylogenetic comparative methods. *Current Biology* **27**, R333–R336 (2017).
9. Ho, L. S. T. *et al.* phylolm: Phylogenetic Linear Regression. (2024).
10. Bartoszek, K. *et al.* Fast MVSLOUCH : Multivariate Ornstein–Uhlenbeck-based models of trait evolution on large phylogenies. *Methods Ecol Evol* **15**, 1507–1515 (2024).
11. Zhang, Y. *et al.* Multivariate Coevolution Shapes Life-History Strategies Across Amniotes. 2025.03.09.642183 Preprint at https://doi.org/10.1101/2025.03.09.642183 (2026).

12. Revell, L. J. phytools: an R package for phylogenetic comparative biology (and other things). *Methods in Ecology and Evolution* **3**, 217–223 (2012).

13. Bartoszek, K. *et al.* Model Selection Performance in Phylogenetic Comparative Methods Under Multivariate Ornstein–Uhlenbeck Models of Trait Evolution. *Syst Biol* **72**, 275–293 (2023).

14. Reitan, T., Schweder, T. & Henderiks, J. Phenotypic evolution studied by layered stochastic differential equations. *Ann. Appl. Stat.* **6**, (2012).
